# Brazil Nut Effect Drives Pattern Formation in Early Mammalian Embryos

**DOI:** 10.1101/2021.02.26.433068

**Authors:** Zheng Guo, Jie Yao, Xu Zheng, Jialing Cao, Zheng Gao, Shuyu Guo, Dandan Qin, Min Tan, Bo Wang, Fanzhe Meng, Jing Zhang, Lei Li, Jing Du, Yubo Fan

## Abstract

The formation of three-dimensional ordered spatial patterns, which is essential for embryonic development, tissue regeneration, and cancer metastasis, is mainly guided by the chemical concentration gradient of morphogens. However, since no chemical concentration gradient has been observed in the early embryonic development (pre-implantation) of mammals, the pattern formation mechanism has been unsolved for a long time. During the second cell fate decision of mouse embryos, the inner cell mass (ICM) segregates into topographically regionalized epiblast (EPI) and primitive endoderm (PrE) layers. Here, we report that the segregation process of PrE/EPI precursors coincides with an emerged periodic expansion-contraction vibration of the blastocyst cavity, which induces phase transition in the ICM compartment to a higher fluidity state and generates directional tissue flows. By experiments and modeling, we demonstrate that the spatial segregation of PrE and EPI precursors is mediated by a “Brazil nut effect”-like viscous segregation mechanism in which PrE precursors with low affinity gradually migrate to the surface of ICM along with the tissue flow, while EPI precursors with high affinity remains inside ICM under cavity vibration. Artificially manipulation of the frequency and amplitude of cavity vibration could control the process of spatial separation as well as lineage specification of PrE/EPI. Furthermore, disruption of the cavity vibration in the initial stage after segregation could reverse the ICM cells back to a mixed state. Therefore, this study reveals a fundamental mechanism that guarantees the robustness of cell segregation and pattern formation without specific morphogens in early mammalian embryos. Our model also emphasizes a conserved function of cavity structure that widely exists in organisms as an energy reservoir and converter between different forms, such as chemical and mechanical energy.

## INTRODUCTION

Spatiotemporally accurate pattern formation is very common in embryonic development. The dorsal-ventral pattern formation in the embryonic development of Drosophila and the pattern formation of neuronal subtypes in the development of vertebrate neural tube are all regulated by the gradient of morphogen^1, 2^. However, in early mammalian embryos composed of several types of cells, there have been no reports of chemical gradients used to form patterns. In early mouse embryos, the first cell lineage specification begins at the 8- to 16-cell stage, resulting in extraembryonic trophectoderm (TE) formation by polar cells enclosing the pluripotent inner cell mass (ICM) composed of apolar cells ^3^. At approximately the 32-cell stage, blastocysts cavity is formed at the interface of the ICM and TE with hyperosmotic fluid secreted by Na^+/^K^+^ ATPases in TE cells ^4–6^. During the development of blastocyst, ICM cells are further segregated into two topographically regionalized lineages, the epiblast (EPI) and primitive endoderm (PrE). EPI gives rise to the whole fetus and extraembryonic mesoderm ^7, 8^, while PrE is a morphologically distinct epithelium separating the epiblast from the blastocyst cavity and is important for the establishment of the yolk sac ^9–11^. Thus, at the time of implantation, the embryo is patterned as a pluripotent epiblast localized beside a fluid lumen and encapsulated by two extraembryonic tissues, the TE and the PrE.At present, the factors that are considered necessary for the plasticity of mammalian embryonic development include geometric constraints, feedback between mechanical and biochemical factors, and cellular heterogeneity^12^.

Progenitors of PrE/EPI lineages can be first identified in early blastocysts by the expression of Nanog in EPI precursors and Gata6 in PrE precursors ^13^. Initially, in the early stages of blastocyst development, EPI precursor cells and PrE precursor cells exist in a mixed “salt and pepper”-like distribution throughout the ICM compartment ^14, 15^. During blastocyst development, EPI and PrE cells are segregated by cell polarity (positional effect) and regulation of transcriptional program in ICM cells ^13–15^. The FGF/ERK signaling pathway has been shown to bias the EPI/PrE fate decision during blastocyst development. EPI precursor cells express FGF4, while PrE precursor cells upregulate FGF receptor 2 (FGFR2) expression ^16, 17^. FGF4 in ICM cells is elevated by Oct4 and Sox2 and binds to FGFR2 in adjacent cells to downregulate Nanog expression and upregulate Gata6 expression ^18–20^. The whole process is accompanied by cell migration in the ICM, indicating a cell sorting process. Differential adhesion-mediated cell sorting processes have been theoretically proposed for the segregation of ICM cells. However, it has not been confirmed in vivo and guarantees only the separation of EPI/PrE precursor cells. The directional signal that drives PrE and EPI precursor cells to allocate to their target positions to complete two-layer pattern formation has remained an open question for a long time.

In this work, combining experiments and modeling, we demonstrate that a “Brazil nut effect”-like viscous segregation mechanism mediate the PrE/EPI cell segregation, pattern formation and maintenance.

## RESULTS

### 1. Topographical regionalization of PrE/EPI layers requires cavity vibration

It has been reported that during the development of blastocysts, PrE precursors move toward the ICM-cavity interface, whereas EPI precursors retain inside the ICM compartment ^21^. In our experiment, we visualized the migration of PrE precursors from stage E3.5 through time-lapse imaging of PrE-specific reporter (*Pdgfra^H2B-GFP/+^*, ^22^) embryos (Figure 1a, b and Supplemental movie S1). By analyzing the relative distance of the GFP-positive cells to the center of the ICM surface ^21^ (Figure 1c), we found that at the initial hours of observation, no significant migration of PrE precursors showed little migration; however, after 8.5 hrs (approximately stage E3.75), the distance of GFP-positive cells to the center of ICM surface was progressively reduced, indicating that the relative allocation of PrE precursors tended to approach the center of the ICM surface (Figure 1d and e). Concurrently, we found that at the beginning of observation, the cavity of the blastocyst continuously expanded, and after E3.75, the blastocoel began to display rhythmic vibration (Figure 1f, g and Supplemental movie S2), which was accompanied by the significant migration of PrE precursors (Figure 1d and e). Interestingly, during rhythmic cavity vibration, each cavity contraction was always coincident with a prominent migration of PrE cells toward the ICM surface (Figure 1e and g). Moreover, accompanied with the emergence of rhythmic cavity vibration, the collective cellular motion in ICM compartment was switched from relatively static state to dynamic state, indicated by cellular motion speed measured by particle image velocimetry (PIV) analysis (Figure 1h, i and Supplemental movie S3). This fluidity transition of ICM was also indicated by assessing the cell shape index p^0^, the median ratio of the perimeter to the square root area of the cells. According to the vertex model, if the cell shape index of the system increases to p*^0^ ≈ 3.81, a transition from a jammed, solid-like state to an unjammed, fluid-like state occurs ^23^. The cell shape index in blastocysts after E3.75 was significantly higher than early blastocysts prior to the emergence of rhythmic cavity vibration (Supplementary Figure S1). These observations indicate a possible correlation between PrE cell movement and cavity vibration.

**Figure 1.**
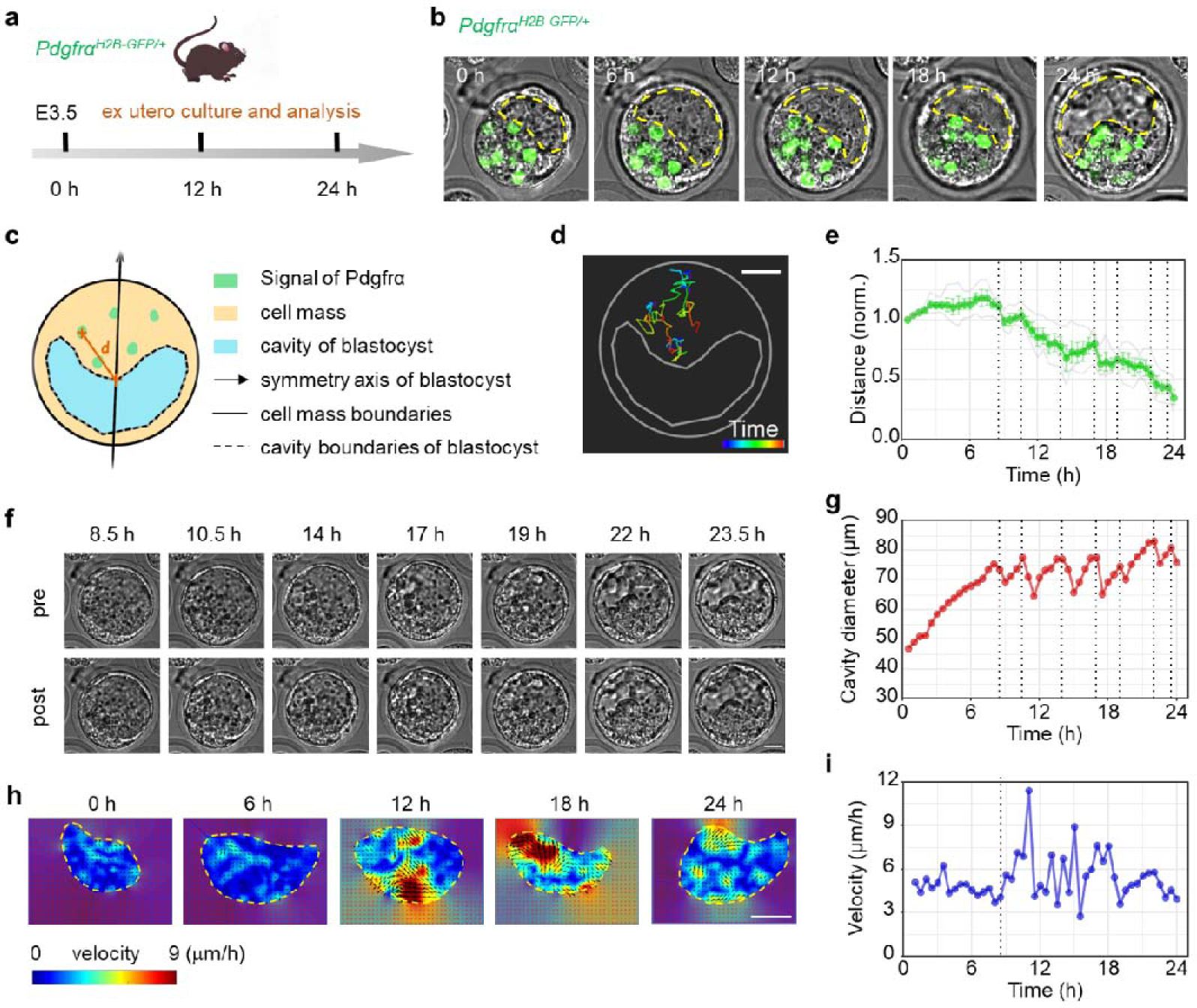
The migration of PrE precursors is consistent with the periodic vibration of the blastocyst cavity. (a) Schematic diagram of in vitro culture of *Pdgfr*α*^H2B-GFP/+^* mouse blastocysts. (b) Time-lapse image of a representative *Pdgfr*α*^H2B-GFP/+^* mouse blastocyst during development. The yellow dotted line denotes the boundary of the cavity. Time (upper left) is shown as hours after E3.5. Scale bar, 20 μm. (c) Schematic diagram of measuring Pdgfrα-GFP positive cells to the center of the ICM surface. Connect the geometric center of the ICM with the geometric center of the cavity to obtain the symmetry axis of the embryo. The intersection of the symmetry axis of the embryo and the boundary of the ICM near the cavity is regarded as the center of ICM surface. Calculate the distance (d) from the center of the cell expressing *Pdgfr*α*^H2B-GFP/+^* to the center of ICM surface. (d) Trajectory of 4 Pdgfrα-GFP positive cells during blastocyst development. Time bar (Lower right), 0 ∼ 24 h. Scale bar, 20 μm. (e) Quantification of the distance of the Pdgfrα-GFP positive cells to the center of the ICM surface over time (thin gray lines are traces of individual cell, green line is traces of the average distance, dotted line indicates the moment when the blastocyst contracted, n = 4). The distance from the Pdgfrα-GFP positive cell at 0 h to the center of the ICM surface is normalized as 1. (f) The representative images of blastocyst prior and post contraction in each period of vibration. Scale bar, 20 μm. (g) Quantification of the blastocyst diameter over time (dotted line indicates the moment when the blastocyst contracted). (h) Time-lapse of the heat map of the velocity magnitude field in ICM during blastocyst development analyzed by PIV. The yellow dotted line denotes the boundary of ICM. Scale bar: 20 μm. (i) The statistical analysis of cell speed (rms velocity) measured by PIV in ICM compartment during blastocyst development. Results in (a) – (i) are obtained in the same representative embryo and repeated for at least three times.

The formation and expansion of the blastocyst cavity stems from the hyperosmotic fluid in the cavity, which generates pressure on the outside TE layer. As the TE layer bears significant tension, rapid cavity contraction occurs when its integrity is disrupted by cell division ^24, 25^. By assessing the tension of embryos indicated by the deformation of TE cells (Supplemental Figure S2a-c) and cavity recoil speed after UV laser cutting (Supplemental Figure S2d and e), we showed that TE tension first increased and then declined during blastocyst development. Thus, We obtained the cortical tension during the development of blastocysts from E3.5 to E4.25 and found that the vibration frequency of the cavity was increased at E3.5 and E4.25 (Supplemental Figure S2f). Meanwhile, significant movement of PrE precursors to the cavity surface occurred at E3.5 and E4.25, when the embryo showed drastic vibration (Supplemental Figure S2g). The cavity vibration and migration of PrE precursors showed a correlation with each other during blastocyst development (Supplemental Figure S2h).

Next, to assess whether the periodic vibration of blastocysts is essential for the directed movement of PrE cells to ICM surface, we suppressed cavity vibration by decreasing hydraulic pressure with hyperosmotic culture medium. The results showed that compared with the control group (cultured in normal KSOM medium), the TE tension and frequency of cavity vibration were both significantly reduced in the hyperosmotic treatment group (Figure 2a and b), and cell motility was significantly inhibited by hypertonic treatment (Figure 2c, d and Supplemental movie S4). As mentioned above, contraction of the blastocyst cavity was generated by cell division in the TE layer to release cortical tension. Embryos were treated with aphidicolin to prevent cell division. As shown in Figure 2e, cavity contraction was reduced by inhibition of cell division. Moreover, ICM fluidity and PrE precursors movement were both significantly suppressed by aphidicolin-treatment (Figure 2f-i). To further evaluate the function of periodic cavity vibration in the spatial segregation of PrE/EPI cells, *Pdgfra^H2B-GFP/+^* embryos were treated with hypertonic medium or normal medium from stage E3.5 (Figure 2j). At the end of 21 hrs of treatment, GFP-positive cells were enriched at the ICM/cavity boundary in the control embryos, while the distribution of GFP-positive cells was still random in the hypertonic group (Figure 2k and l). Then, the embryos in hyperosmotic medium were washed out and moved to KSOM medium for continuous culture (Figure 2j). After 24 hrs, GFP-positive cells were layered at the boundary of the ICM and blastocyst cavity in both groups (Figure 2k and l), indicating that interruption of cavity vibration prevented the segregation of PrE/EPI. These results suggest that the periodic vibration of cavity is essential for the topographical regionalization of PrE/EPI layers.

**Figure 2.**
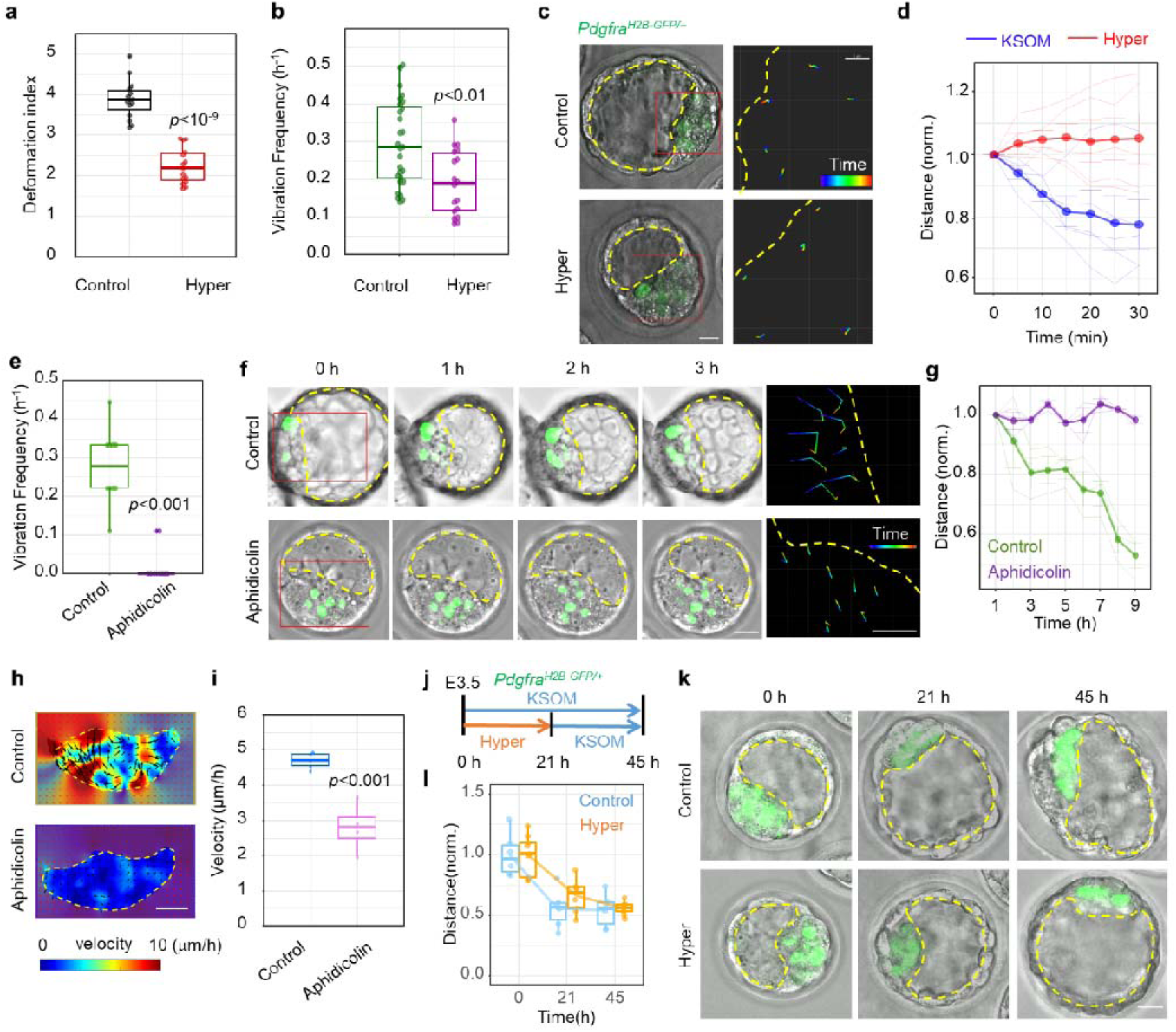
Inhibition of cavity vibration prevents the topographical regionalization of PrE. (a) Deformation index (aspect ratio) of TE cells in control (n = 15 cells from 5 embryos) and hypertonic medium treated (n = 15 cells from 5 embryos) embryos. (b) Vibration frequency of blastocyst cavity in control (n= 28 embryos) and hypertonic medium treated (n = 18 embryos) embryos. (c) Movement trajectory of Pdgfrα-GFP positive cells in control and hypertonic medium treated embryos. The yellow dotted line denotes the boundary of the cavity. The image on the right is a magnification of the red frame area of the left image. Scale bar: left, 25 μm, right, 5 μm. Time bar (lower right), 0 ∼ 30 min. (d) Quantification of the distance of the Pdgfrα-GFP positive cells to the center of the ICM surface over time in control and hypertonic medium treated embryos (thin lines are traces of individual cell, thick line is traces of the average of cells, dots are averages, n = 5 cells). The distance from the Pdgfrα-GFP positive cell at 0 h to the center of the ICM surface is normalized as 1. (e) Vibration frequency of blastocyst cavity in control and phidicolin-treated embryos (n = 10 embryos). (f) Time-lapse (left, scale bar, 20 μm) and movement trajectory (right, scale bar, 5 μm. Time bar, 0 ∼ 3 h.) of Pdgfrα-GFP positive cells in control and aphidicolin-treated embryos. The yellow dotted line denotes the boundary of the cavity. The image on the right is a magnification of the red frame area of the left image. (g) Quantification of the distance of the Pdgfrα-GFP positive cells of the control (green, n = 9 cells from 3 embryos) and the aphidicolin treatment (purple, n = 9 cells from 3 embryos) to the center of the ICM surface over time (thin lines are traces of individual embryo, thick line is traces of the average of embryos). The distance from the Pdgfrα-GFP positive cell at 1 h to the center of the ICM surface is normalized as 1. (h) The representative heat map of the velocity field in ICM of control and aphidicolin-treated embryos analyzed by PIV. The yellow dotted line denotes the boundary of ICM. Scale bar, 20 μm. (i) The statistical analysis of cell speed (rms velocity) of ICM compartment measured by PIV in control (n = 5 embryos) and aphidicolin-treated embryos (n = 4 embryos). The distance from the Pdgfrα-GFP positive cell at 0 h to the center of the ICM surface is normalized as 1. (j) Schematic diagram of the process of hypertonic treatment of blastocysts. The length of the arrow represents the duration of embryo culture. (k) Time-lapse images of Pdgfrα-GFP embryos in control and hypertonic treatment. The yellow dotted line denotes the boundary of the cavity. Scale bar, 20 μm. (l) Quantification of the distance of the Pdgfrα-GFP positive cells to the center of the ICM surface over time in the control (n = 6 cells from 2 embryos) and the hypertonic medium treated (n = 9 cells from 3 embryos) embryos. The experiments were repeated for at least three times.

### 2. Cavity vibration induces PrE/EPI pattern formation through viscous segregation

Next, we sought to determine the mechanism underlying the promotion of PrE/EPI segregation by cavity vibration. As mentioned above, cavity vibration elevated the fluidity of ICM compartment (Figure 1h, i and Figure 2h, i). Further study showed that during a period of contraction-expansion vibration, converged tissue flows from the TE side toward the center of the ICM/cavity boundary was observed in the ICM compartment after each contraction. Moreover, the direction of the tissue flows were coincident with the movement of PrE cells (Figure 3b and Supplemental movie S5). Meanwhile, during the expansion of blastocoels, significant tissue flows was observed at the ICM surface parallel to the ICM/cavity boundary, which was accompanied by the lateral movement of PrE cells (Figure 3c, d and Supplemental movie S6). By mechanical analysis and simulation, we demonstrated that the dynamic tissue flows stemmed from the contraction or expansion of the embryo (Figure 3e, Supplemental movie S7, and Mechanical Model in the Supplemental materials). The frequency and amplitude of the blastocyst vibration are equivalent to power. When the power of the blastocyst vibration reaches a certain threshold, the cell can use the energy to perform directional movement. During the contraction phase of the blastocyst, the adhesion between cells can be broken, so that the movement of the cells is remarkable. However, during the expansion phase of the blastocyst, the adhesion between cells cannot be broken, so the position of the cells is relatively unchanged. Cell mucus is a non-Newtonian fluid, and it will become thinner and easier to flow when subjected to repeated shearing forces. Since the time scale of the contraction phase was much shorter than that of the expansion phase, through several repetitions of the asymmetric slow-expansion/rapid-contraction period, ratchet effect would be introduced that led to symmetry-breaking and directed cell migration ^26^. The ratchet effect can provide directed transport of small particles in the absence of any significant macroscopic driving force, and has been found important in many microscale systems ^26^. Here this ratchet mechanism served to establish the background tissue flow, which resulted in directed cell migration towards the center of the blastocyst during several expansion/contraction periods. On the other hand, numeric evidences have shown that during the segregation of EPI and PrE precursors, these two types of cells showed differences in cohesive property and motility. Compared with EPI precursors, PrE precursors possess lower cohesiveness and higher mobility ^27–33^. We also found that, during blastocyst development, an increased cell aggregation tendency was observed in EPI cells compared with PrE cells (Figure 3f). Due to the unequal interval and transport ability between slow expansion and rapid contraction, the PrE cells with low cohesiveness and high mobility can be driven towards the center of the blastocyst. This mechanism is analogous to the viscous segregation of mixed granules based on different adhesive properties under exogenous energy input, for example, the “Brazil nut effect”. The Brazil nut effect means that if a pile of nuts of different sizes or frictions is mixed into a container and then an additional vibration is applied, the largest Brazil nut will usually rise to the surface, while the smaller other nuts will settle to the bottom, which has important implications for many industrial processes and geophysical processes^34–37^. In our system, the vibration of the cavity provides an exogenous energy input as disturbance and caused phase transition with increased tissue flow in the ICM compartment, which gave rise to the segregation of PrE and EPI cells depending on their distinct cohesivenes. This hypothesis could be supported by a transient intermediate state which was observed during PrE/EPI segregation before pattern formation completion, in which Pdgfrα-positive cells were enriched as an “inverted pyramid” at the central region of the ICM/cavity boundary and embraced by Nanog-positive cells (Figure 3g and h).

**Figure 3.**
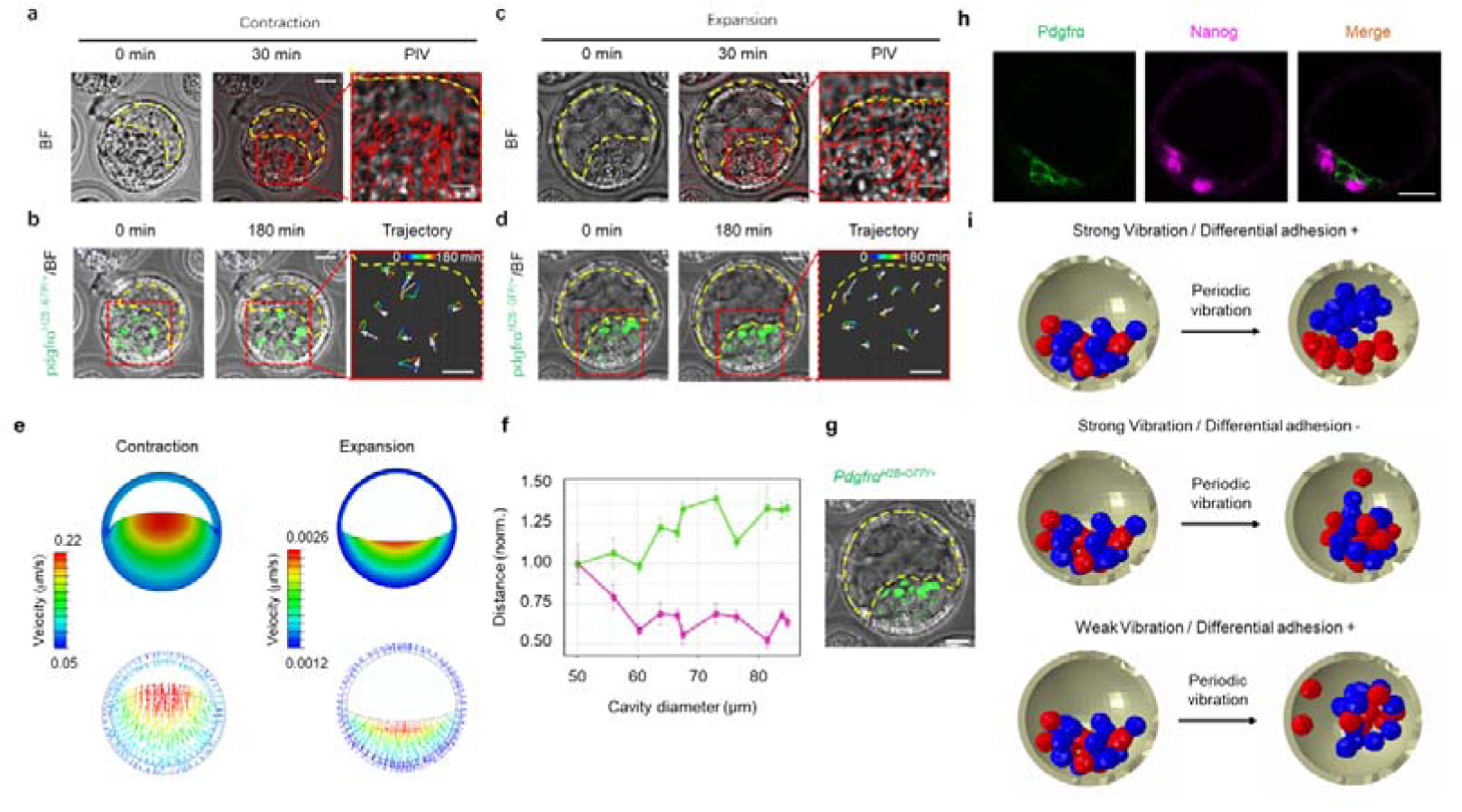
Cavity vibration induces dynamic tissue flows in ICM to drive cell movement. (a and c) The velocity field of ICM compartment during blastocyst contraction (a) and expansion (c) analyzed by PIV. The length and direction of the arrow indicate the magnitude and direction of the speed at that point, respectively. The image on the right is a magnification of the red frame area of the left image. The yellow dotted line denotes the boundary of the cavity. (b and d) Movement trajectory of Pdgfrα-GFP positive cells during contraction (b) and expansion (d) of blastocyst. The image on the right is a magnification of the red frame area of the left image. The yellow dotted line denotes the boundary of the cavity. The white arrow indicates the displacement of the cell. Scale bar, 5 μm. Time bar (upper right), 0 ∼ 180 min. (e) A simulation of motion pattern of embryo during contraction and expansion with a simplified finite element model. Images selected to show the distribution of velocity magnitude (top) and orientation (bottom) at the 60 s of contraction (left) and 3600 s of expansion (right). (f) Quantification of the distance of the Nanog^+^/Nanog^+^ cells (magenta) and Pdgfrα^+^/Pdgfrα^+^ cells (green) in blastocysts with different cavity diameters (n = 11 embryos). The average value of the distance from the Pdgfrα-GFP positive cells of the embryo with a diameter of 50μm to the center of the ICM surface is normalized as 1. (g) Representative *Pdgfr*α*^H2B-GFP/+^* embryo image (right) of the Pdgfrα-GFP positive cells enriched as an “inverted pyramid” at the central region of ICM/cavity boundary. The yellow dotted line denotes the boundary of the cavity. Scale bar, 20 μm. (h) Representative immunofluorescence images of Pdgfrα and Nanog in embryo at “inverted pyramid” intermediate state. Scale bar, 25 μm. The experiments were repeated for at least three times. (i) A Simulations of cell viscous segregation during cavity vibration with a simplified finite element model. The model was to illustrate the effect of cohesive properties and periodic contraction-expansion of embryos on the separation of PrE and EPI cells. Top: The separation process of PrE and EPI cells with different cohesive properties (EPI: red, strong cohesiveness; PrE: blue, weak cohesiveness) under periodic embryo contraction-expansion. Middle: The separation failed when cohesive properties of cells were the same. Bottom: The separation failed when embryo contraction speed reduced to one-third.

To evaluate this Brazil nut effect hypothesis, we monitored the migration of chemically inert fluorescent microbeads that were injected into the ICM compartment of early blastocysts (Supplemental Figure S3a). The results showed that although the microbeads were initially randomly distributed in the ICM compartment, they were enriched at the interface of the ICM/cavity in most embryos after several hours (Supplemental Figure S3b and c). Directed movement of microbeads was also observed after embryo contraction (Supplemental Figure S3d and Supplemental movie S8). Next, we conducted computational modeling using a finite element simulation and found that the allocation and sorting of PrE/EPI precursors during the slow-expansion/rapid-contraction periods could be recapitulated by the segregation of cohesive granular flow driven by asymmetry vibration (Figure 3i, Supplemental Figure S4a, Supplemental movie S9 and Mechanical model in the Supplemental materials). Moreover, the simulation results predicted that successful segregation of cells relies on the difference in cell cohesive properties and the asymmetry of expansion-contraction speed (Figure 3i).

On the other hand, when the embryo was in the “inverted pyramid” intermediate state, position effect (e.g. polarity signals) together with the lateral tissue flows at the surface of the ICM during the expansion of the blastocyst cavity could drive the rearrangement of Pdgfrα-positive cells to line as a monolayer. In addition, we found that significant F-actin bundles were enriched at the boundary of Pdgfrα-positive cells and the cavity in the “inverted pyramid” intermediate state (Supplemental Figure S4b), and the outermost Pdgfrα-positive cells displayed a “Rose-like” distribution at the surface of the ICM (Supplemental Figure S4c), indicating the contribution of actomyosin contractility in the epithelialization of PrE cells to complete the pattern formation of ICM.

### 3. PrE/EPI spatial segregation and lineage specification could be controlled by cavity vibration

To further investigate the function of cavity vibration in the lineage specification of PrE/EPI, we interrupted blastocoel tension by different chemical and physical treatments. As mentioned above, hypertonicity treatment reduced TE tension (Figure 2a) and suppressed cavity vibration (Figure 2b) as well as the migration of Pdgfrα-positive cells (Figure 2c, d and j-l). We further confirmed that hypertonic treatment of embryos from stage E3.5 for 24 hrs significantly suppressed the spatial segregation of PrE/EPI precursors compared with control embryos (Figure 4a and b). Moreover, the lineage specification of PrE/EPI cells was also interrupted by hypotonic treatment (Figure 4a and c-e). Since the hyperosmotic solution secreted into the blastocyst cavity is mediated by the Na^+^/H^+^ exchanger isoform NHE-3 in TE cells, we treated embryos with the NHE3 inhibitor S3226, which has been shown to inhibit the expansion of blastocysts ^38^. The vibration frequency of the blastocyst cavity in the S3226 group was significantly reduced compared with that of the control group (Supplemental Figure S5a). The results showed that the spatial segregation and lineage specification were both inhibited by S3226 treatment (Supplemental Figure S5b-f). It has been demonstrated that the expansion and contraction of the blastocyst cavity requires cell contractility by the actomyosin cytoskeleton ^25^. Thus, we treated embryos with the actomyosin inhibitors blebbistatin (Bleb) and cytochalasin A (CA) to inhibit cavity vibration (Supplemental Figure S6a and S7a). Both of these actomyosin inhibitor treatments dramatically disrupted the spatial segregation of PrE/EPI cells and prevented their lineage specification (Supplemental Figure S6 and S7). Together, these data show that interruption of cavity vibration inhibits PrE/EPI spatial segregation and lineage specification.

**Figure 4.**
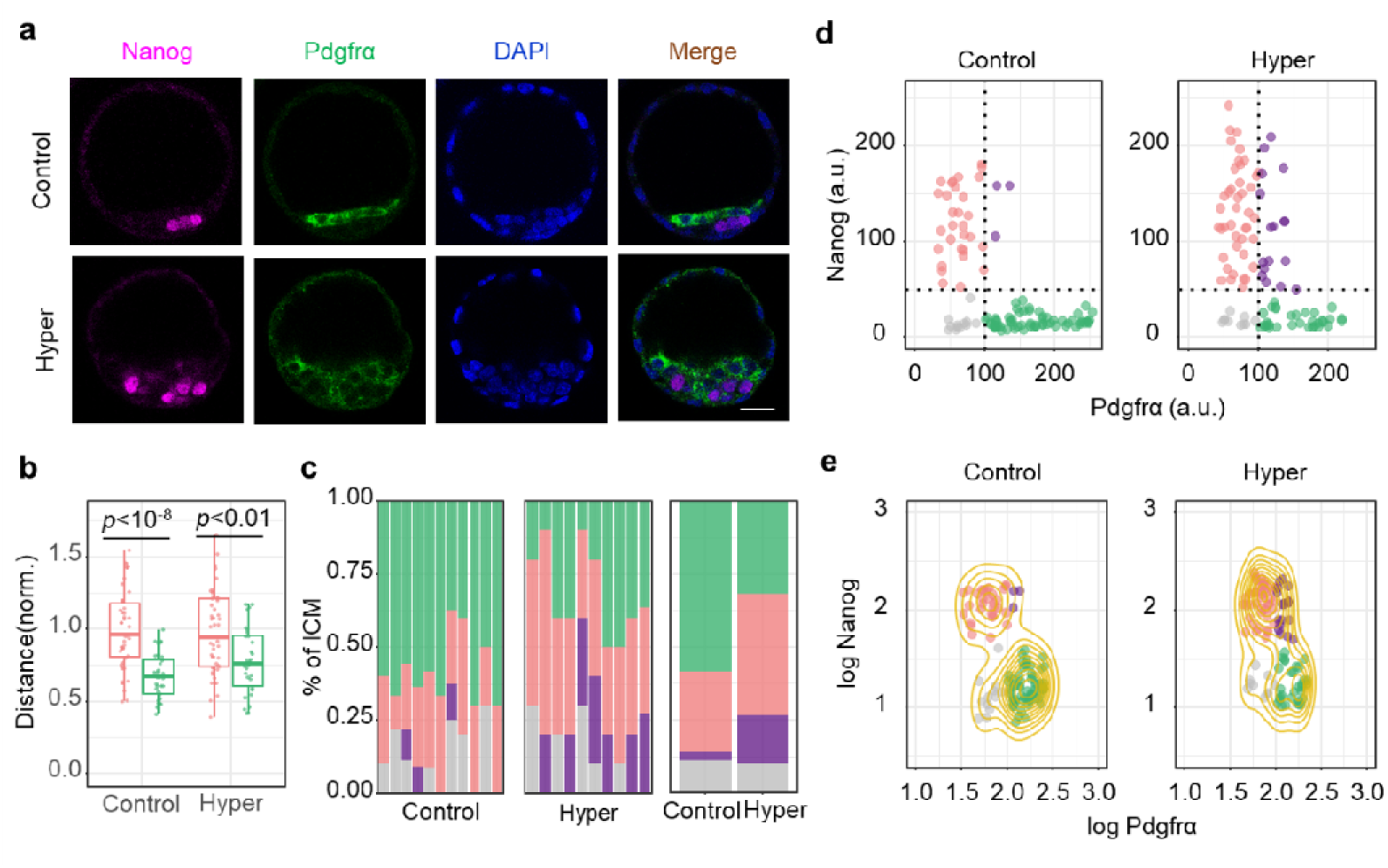
Hypertonic treatment inhibits the spatial segregation and lineage specification of PrE/EPI. (a) Representative immunofluorescence images of the E3.5 embryos fixed after 24 hours of culture in hypertonic (hyper) or normal medium (control). Scale bar, 25 μm. (b) The distance of Nanog and Pdgfrα positive cells to the center of the ICM surface in control (Nanog: n = 46 cells from 9 embryos, Pdgfrα: n = 33 cells from 9 embryos) and hypertonic treatment (Nanog: n = 49 cells from 7 embryos, Pdgfrα: n = 30 cells from 7 embryos). The mean value of the distance from the nanog cells of the control group to the center of the ICM surface is normalized as 1. (c) Average ICM composition at the end of the culture period in normal or hypertonic medium, shown as % of the ICM. Double negative DN, gray, (Nanog-, Pdgfrα-); Double positive DP, purple, (Nanog+, Pdgfrα+); EPI, red, (Nanog+, Pdgfrα-); PrE, green, (Nanog-, Pdgfrα-). Control: n = 108 cells from 11 embryos. Hypertonic: n = 101 cells from 10 embryos. (d) Scatter plot of fluorescence intensity levels of Nanog and Pdgfrα after hypertonic treatment. (e) Scatter plots for same data as in (d), represented as logarithm. The yellow contour lines show the density. The experiments were repeated for at least three times.

Next, we wondered whether the segregation of PrE/EPI cells could be facilitated by reinforced cavity vibration. We first treated E3.25 stage embryos with hypotonic medium, which significantly increased the frequency of cavity vibration (Figure 5a). At the end of treatment for 24 hrs, the spatial segregation and lineage specification of EPI/PrE cells were almost complete in the hypotonic group, while these two types of cells were still mixed in the control embryos (Figure 5b-f). This phenomenon was confirmed by lysophosphatidic acid (LPA) treatment, which could promote the activity of NHE-3 ^39^. The treatment of LPA also enhanced cavity vibration (Supplemental Figure S8a) and accelerated the spatial segregation and lineage specification of PrE/EPI cells EPI/PrE segregation (Supplemental Figure S8b-f). In summary, these results indicate that cavity vibration plays a pivotal role in the spatial segregation and fate decision of PrE/EPI.

**Figure 5.**
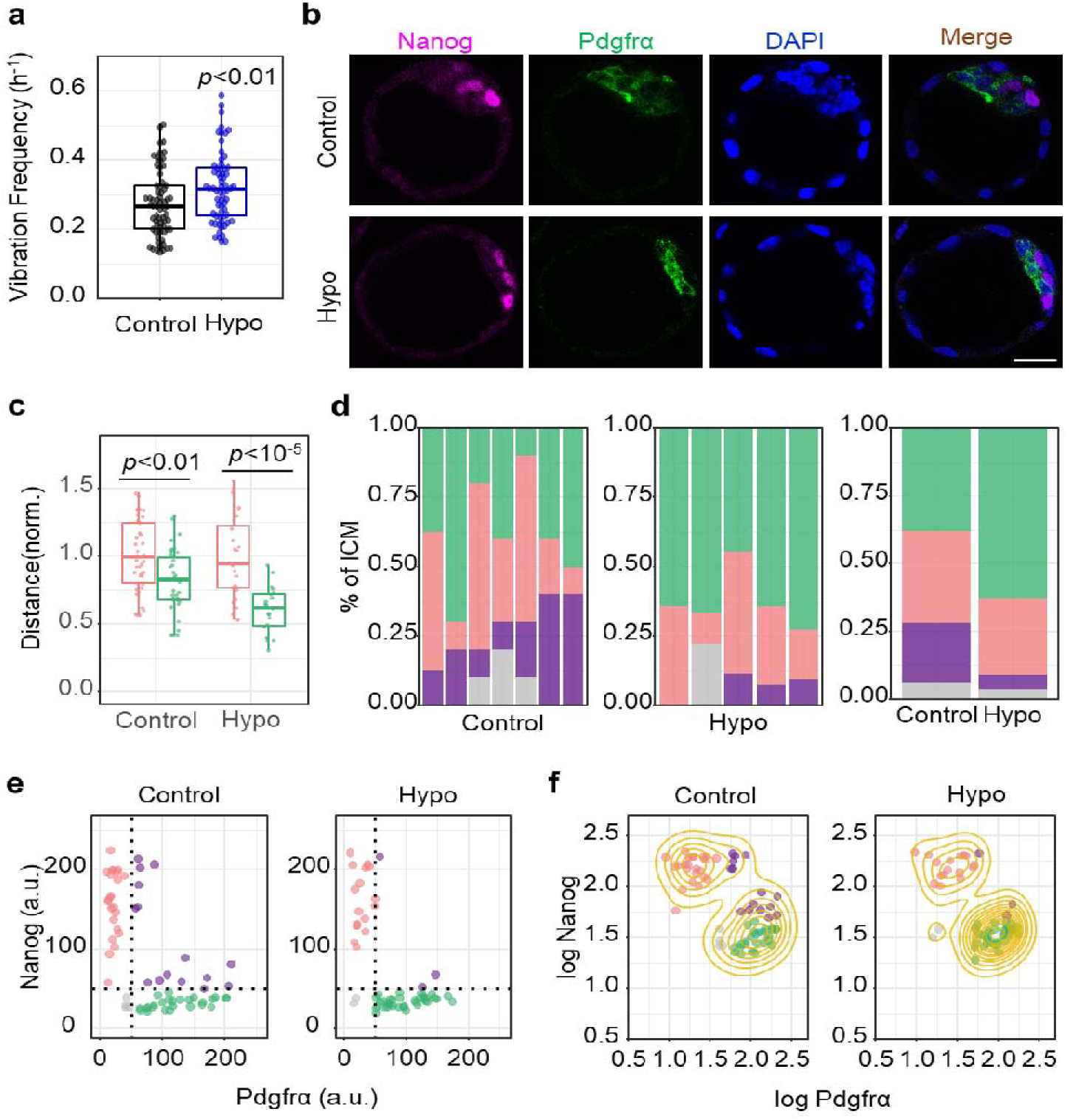
Hypotonic treatment promotes the spatial segregation and lineage specification of PrE/EPI. (a) Vibration frequency of blastocyst cavity in control (n = 66 embryos) and hypotonic-treated (Hypo, n = 64 embryos) embryos. (b) Representative immunofluorescence images of the E3.25 embryos fixed after 24 hours of culture in hypotonic or normal medium. Scale bar, 25 μm. (c) The distance of Nanog and Pdgfrα positive cells to the center of the ICM surface in control (Nanog: n = 41 cells from 8 embryos, Pdgfrα: n = 35 cells from 8 embryos) and hypotonic treated (Nanog: n = 24 cells from 5 embryos, Pdgfrα n = 21 cells from 5 embryos) embryos. The mean value of the distance from the nanog cells of the control group to the center of the ICM surface is normalized as 1. (d) Average ICM composition at the end of the culture period in embryos treated with normal or hypotonic medium, shown as % of the ICM. (e) Scatter plot of fluorescence intensity levels of Nanog and Pdgfrα after hypotonic treatment. Double negative, DN, gray, (Nanog-, Pdgfrα-); Double positive, DP, purple, (Nanog+, Pdgfrα+); EPI, red, (Nanog+, Pdgfrα-); PrE, green, (Nanog-, Pdgfrα-). Control: n = 68 cells from 7 embryos. Hypotonic: n = 57 cells from 5 embryos. (f) Scatter plots for same data as in (e), represented as logarithm. The yellow contour lines show the density. The experiments were repeated for at least three times.

### 4. PrE/EPI spatial segregation and lineage specification maintenance requires cavity vibration during the initial stage after two-layer pattern formation

We noticed that the cavity vibration frequency remained at a high level at late blastocyst after pattern formation of PrE/EPI layers (E4.25 in Supplemental Figure S2f). To determine whether the maintenance of ICM regionalization also requires vibration of the cavity, we monitored the effect of hypertonic treatment on the cell allocation dynamics in *Pdgfra^H2B-GFP/+^* blastocysts at different stages. We found that the segregation of PrE/EPI cells could be reversed when hypertonicity was applied at the initial stage after PrE/EPI (within ∼3 hrs after the “inverted pyramid” intermediate stage). However, when hypertonicity was treated to embryos several hours (at least ∼5 hrs) post the intermediate stage, the two-layer pattern could not be reversed (Figure 6a and b), indicating that the vibration of the cavity is required for the maintenance of the PrE/EPI regionalization at the initial stage after segregation.

**Figure 6.**
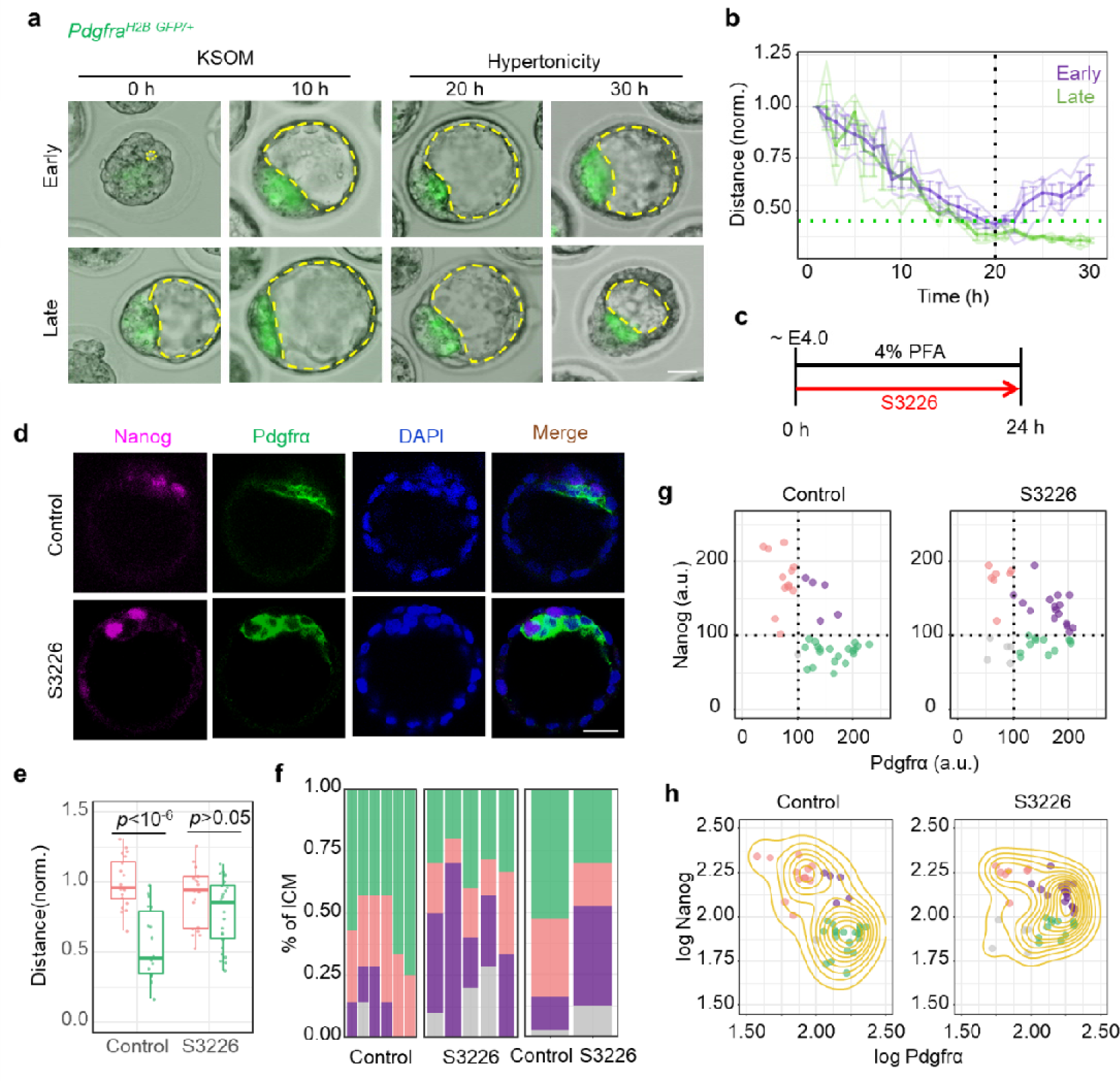
PrE/EPI spatial segregation and lineage specification maintenance requires cavity vibration during the initial stage after two-layer pattern formation. (a) Representative time-lapse images of blastocysts expressing Pdgfrα-GFP in control and hypertonic medium at early or late stage as described in the main text. The yellow dotted line denotes the boundary of the cavity. Scale bar, 25 μm. (b) Quantification of the distance of the Pdgfrα-GFP positive cells to the center of the ICM surface over time in embryos treated with hypertonicity at early (n = 9 cells from 3 embryos) or late(n = 9 cells from 3 embryos) stage. Thin lines are traces of individual embryo, thick line is traces of the average of embryos. The distance from the Pdgfrα-GFP positive cell at 1 h to the center of the ICM surface is normalized as 1. (c) Schematic diagram of the process of S3226 treatment of blastocysts. The length of the arrow represents the duration of embryo culture. (d) Representative immunofluorescence images of the E4.0 embryos treated with S3226 as (c). Scale bar, 25μm. (e) The distance of Nanog and Pdgfrα positive cells to the center of the ICM surface in control (Nanog: n = 20 cells from 4 embryos, Pdgfrα: n = 18 cells from 4 embryos) or S3226-treated (Nanog: n = 19 cells from 4 embryos, Pdgfrα: n = 25 cells from 4 embryos) embryos. The mean value of the distance from the nanog cells of the control group to the center of the ICM surface is normalized as 1. (f) Average ICM composition at the end of the culture period in control or S3226-treated embryos, shown as % of the ICM. Comtrol: n = 38 cells from 6 embryos. S3226: n = 40 cells from 5 embryos. (g) Scatter plot of fluorescence intensity levels of Nanog and Pdgfrα in control or S3226-treated embryos. DN, gray, double negative (Nanog-, Pdgfrα-); DP, purple, double positive (Nanog+, Pdgfrα+); EPI, red, (Nanog+, Pdgfrα-); PrE, green, (Nanog-, Pdgfrα-). (h) Scatter plots for same data as in (g), represented as logarithm. The yellow contour lines show the density. The experiments were repeated for at least three times.

To further determine the reversibility of lineage specification of PrE/EPI by vibration inhibition, Embryos at E4.0 were treated with the NHE-3 inhibitor S3226 for 24 hrs and compared with control embryos fixed at E4.0 after collection (Figure 6c). While most of the control embryos had completed PrE/EPI spatial segregation, most of S3226-treated embryos showed a mixed distribution of cells expressing Nanog or Pdgfrα (Figure 6d and e). Moreover, the lineage specification of PrE/EPI was also interrupted by S3226-treatment (Figure 6f-h). Similar results were observed in embryos treated with hypertonic medium (Supplemental Figure S9). These results suggest that during the initial stage after PrE/EPI two-layer pattern formation, the spatial segregation and lineage specification are both sustained partially by cavity vibration.

### 5. Yap signal is involved in periodic embryo vibration to regulate gene expression of PrE and EPI precursors

To determine the mechanism underlying the promotion effect of cavity vibration in PrE/EPI lineage specification, we analyzed the gene expression level during segregation process. We found that in *Pdgfra^H2B-GFP/+^* embryos, the GFP expression intensity was progressively enhanced after the beginning of cavity vibration (8.5 hr) during the migration of Pdgfrα-positive cells (Figure 7a–c and in the Supplemental movie S10), indicating a positive correlation of gene expression of PrE marker gene and cavity vibration. Moreover, the expression level of Pdgfrα was significantly reduced by hypertonic treatment (Figure 7d) and increased by hypotonic medium (Figure 7e), whereas Nanog expression was little affected by these treatments. Moreover, the ratio of Nanog-positive to Pdgfrα-positive cells was also significantly induced by hypertonicity and reduced by hypotonicity (Figure 7f). Similar results were observed in embryos treated with NHE-3 inhibitor, LPA, blebbistatin, and CA (Supplemental Figure S10). These results indicate that cavity vibration contributes to the gene expression of PrE/EPI precursors during spatial segregation.

**Figure 7.**
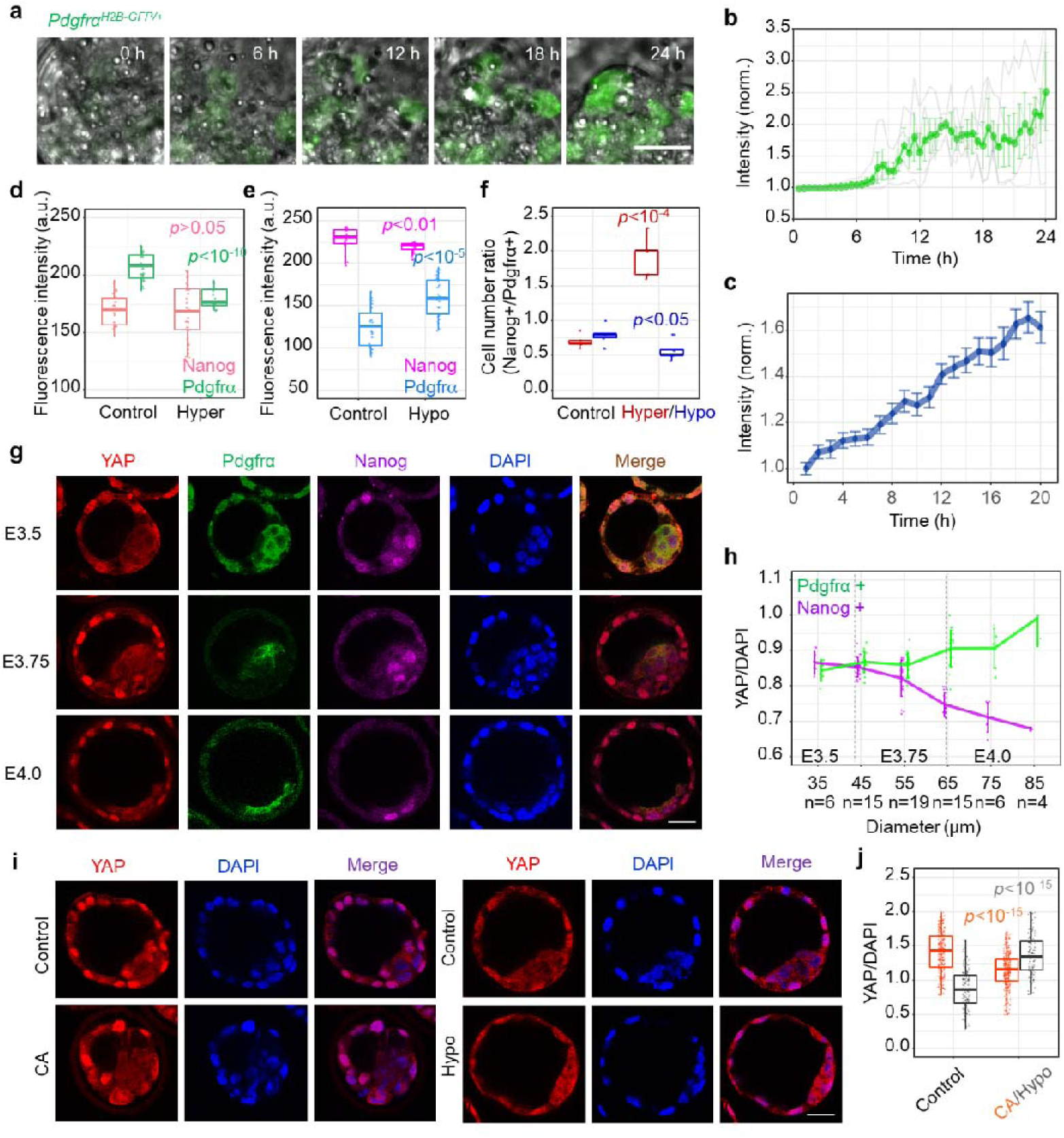
Yap signal is involved in periodic embryo vibration to regulate gene expression of PrE and EPI precursors. (a) Representative time-lapse images of the ICM in a Pdgfrα-GFP expressing embryo during development. Scale bar, 20 μm. (b) Quantification of the fluorescence intensity of Pdgfrα-GFP positive cells in a representative single embryo over time (n = 4 Pdgfrα-GFP positive cells, thin lines are traces of individual cell, thick line is traces of the average of cells). The fluorescence intensity of the Pdgfrα-GFP positive cells at 0h is normalized as 1. (c) Quantification of the fluorescence intensity of Pdgfrα-GFP positive cells in multiple embryos over time. n = 13 embryos. The mean value of the fluorescence intensity of the Pdgfrα-GFP positive cells at 1 h is normalized as 1. (d) The fluorescence intensity of control (Nanog: n = 22 cells from 5 embryos, Pdgfrα: n = 31 cells from 5 embryos) and hypertonicity-treated (Nanog: n = 34 cells from 5 embryos, Pdgfrα: n = 18 cells from 5 embryos) embryos. (e) The fluorescence intensity of control (Nanog: n = 18 cells from 5 embryos, Pdgfrα: n = 23 cells from 5 embryos) and hypotonicity-treated (Nanog: n = 18 cells from 5 embryos, Pdgfrα: n = 33cells from 5 embryos) embryos. (f) The cell number ratio of Nanog^+^/Pdgfrα^+^ in control, hypertonicity or hypotonicity treated embryos. (g) Representative immunofluorescence images of blastocysts at different stages. Scale bar, 25 μm. (h) Quantification of the YAP/DAPI fluorescence intensity in Nanog+ cells and Pdgfrα+ cells in embryos different cavity diameters. The DAPI fluorescence intensity of the nucleus is normalized as 1. (i) Representative immunofluorescence images of E3.5 embryos treated with cytochalasin A (CA) or hypotonicity. Scale bar, 25 μm. (j) The statistical analysis of YAP/DAPI fluorescence intensity of ICM cells in embryos treated with CA (Control: n = 304 cells from 17embryos, CA: n = 304 cells from 15 embryos) or hypotonicity (Control: n = 147 cells from 8 embryos, Hypotonicity n = 108 cells from 7 embryos). The experiments were repeated for at least three times. The DAPI fluorescence intensity of the nucleus is normalized as 1.

In the first lineage specification of mouse embryo, the formation of ICM/TE pattern due to the asymmetric activation of Yes-associated protein (YAP), which has been demonstrated to be a conserved mechanical sensor and is involved in diverse developmental processes ^40^. There is evidence that YAP signaling increases the expression of FGF receptors (FGFRs) in embryonic neural stem cells and lung cancer samples ^41, 42^. Thus, we assessed the potential role of YAP in the lineage specification of PrE/EPI promoted by cavity vibration. During the development of blastocysts, the levels of nuclear YAP in PrE and EPI precursors were comparable at E3.5, however, significant difference in the nuclear YAP level in PrE and EPI precursors was observed after E3.75(Figure 7g and h). Moreover, the nuclear/cytoplasmic ratio of YAP showed an increasing tendency in PrE precursors, while this ratio was gradually reduced in EPI cells (Figure 7g and h). Interruption of cavity vibration by CA treatment significantly suppressed the nuclear level of YAP, while promoted vibration by hypotonic medium elevated the nuclear level of YAP (Figure 7i and j). Since the lineage specification of PrE/EPI is completed at late E4.0, these observations indicate a possible role of YAP in the regulation of PrE/EPI cell fate by cavity vibration. To evaluate this hypothesis, YAP activity was reduced by its inhibitor Verteporfin (VP) during the development of blastocyst. In order to reduce the influence of VP on TE cells, VP treatment was performed at the blastocyst stage after the first fate decision was completed.As shown in Figure 8, VP treatment significantly disrupted the spatial segregation and lineage segregation of PrE/EPI layers. These results indicate that the yap signal is involved in periodic embryo vibration to regulate gene expression of PrE and EPI precursors.

**Figure 8.**
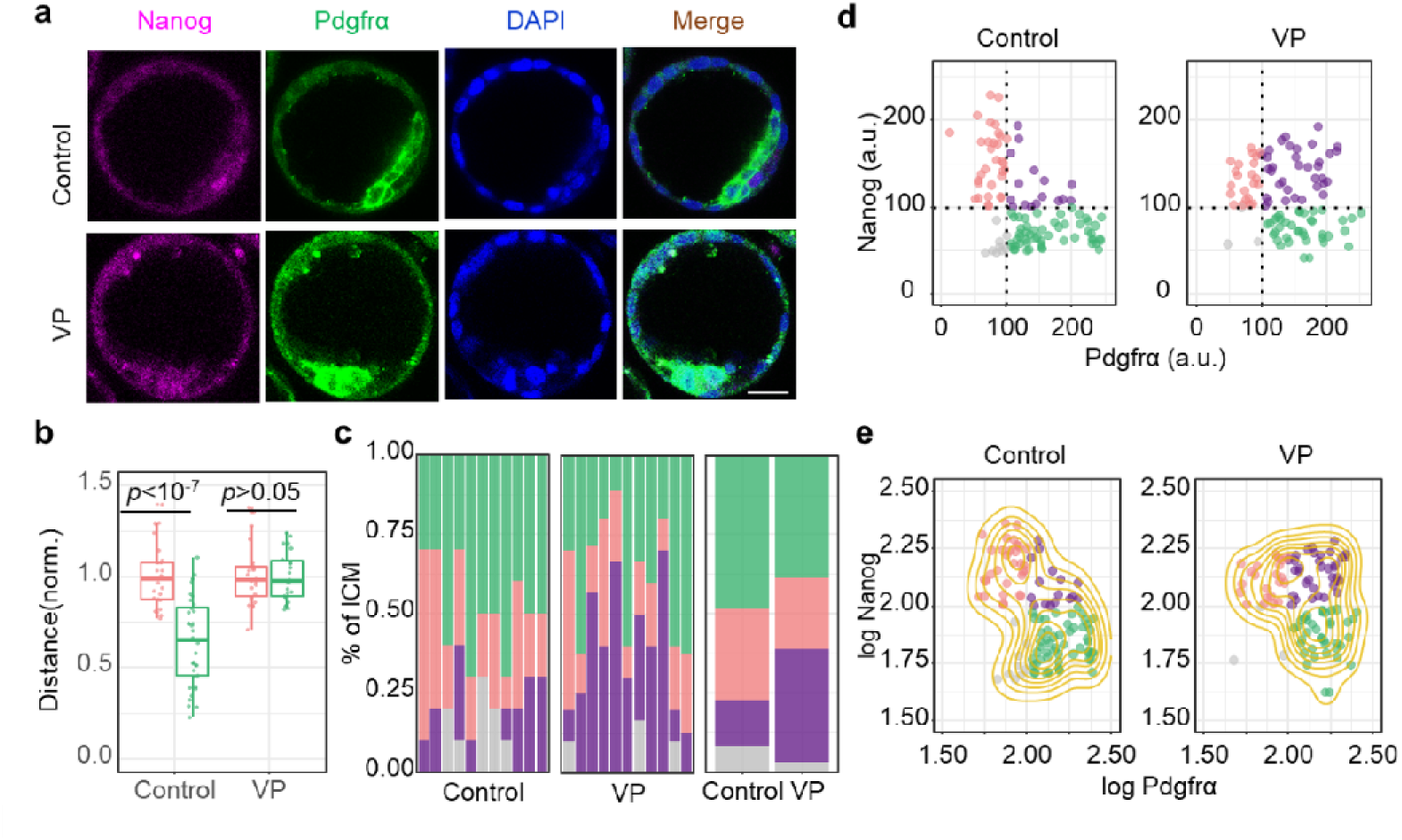
YAP inhibition prevents the spatial segregation and lineage specification of PrE/EPI. (a) Representative immunofluorescence images of the E3.5 embryos fixed after 24 hours of verteporfin (VP) treatment. Scale bar, 25 μm. (b) The distance of Nanog and Pdgfrα positive cells to the center of the ICM surface in control (Nanog: n = 27 cells from 7 embryos, Pdgfrα: n = 36 cells from 7 embryos) or VP treated (Nanog: n = 23 cells from 5 embryos, Pdgfrα: n = 23 cells from 5 embryos) embryos. The mean value of the distance from the nanog cells of the control group to the center of the ICM surface is normalized as 1. (c) Average ICM composition at the end of the culture period in control or VP-treated embryos, shown as % of the ICM. Comtrol: n = 110 cells from 11 embryos. VP: n = 98 cells from 11 embryos. (d) Scatter plot of fluorescence intensity levels of Nanog and Pdgfrα after VP treatment. DN, gray, double negative (Nanog-, Pdgfrα-); DP, purple, double positive (Nanog +, Pdgfrα+); EPI, red, (Nanog +, Pdgfrα-); PrE, green, (Nanog-, Pdgfrα-). (e) Scatter plots for same data as in (d), represented as logarithm. The yellow contour lines show the density. The experiments were repeated for at least three times.

## DISCUSSION

In the development of many animal embryos, morphogens are involved in cell fate determination. For example, in the development of vertebrate neural tube, different neuron subtypes are distributed in precise spatial order according to the concentration gradient of morphogen released by progenitor cells^2^. However, a recent study shows that even in the classic model of morphogen-driven pattern formation, physical factors like different adhesion properties of cells, are essential for this process ^43^. There are few mechanisms that can replace morphogens to drive cell spatial distribution. There is an alternative morphogen mechanism in Dictyostelium discoideum, which is based on selecting cells that initially differentiate in a mixture of salt and pepper and then physically move to produce a coherent tissue ^44^. In early mouse embryos, there is no morphogen gradient has been reported for cell sorting. A computational model of cell sorting indicated that the asymmetric affinity between EPI and PrE cells is not enough to complete spatial isolation in mouse embryos ^30, 31^. Our results show that in addition to the differential affinity between EPI and PrE cells, the mechanical vibration of the cavity is required to complete the segregation of the spatial location. Cavity vibration induces fluidization of ICM and generates directional tissue flows to direct the segregation of PrE/EPI precursors. Tissue flow is common in the process of embryo development and organ formation and is often accompanied by the process of cell migration and cell sorting ^45^. In our model, the asymmetric contraction/expansion speed generates net tissue flows converging toward the surface of the ICM. Under this mechanical force, the low affinity cells move toward the surface and the high affinity cells maintain in the interior region. After arriving at the ICM surface as a “inverted pyramid”, PrE cells finally form a monolayer epithelium near the blastocyst cavity under the action of positional induction ^46^ and actomyosin contraction (Figure 9).

**Figure 9.**
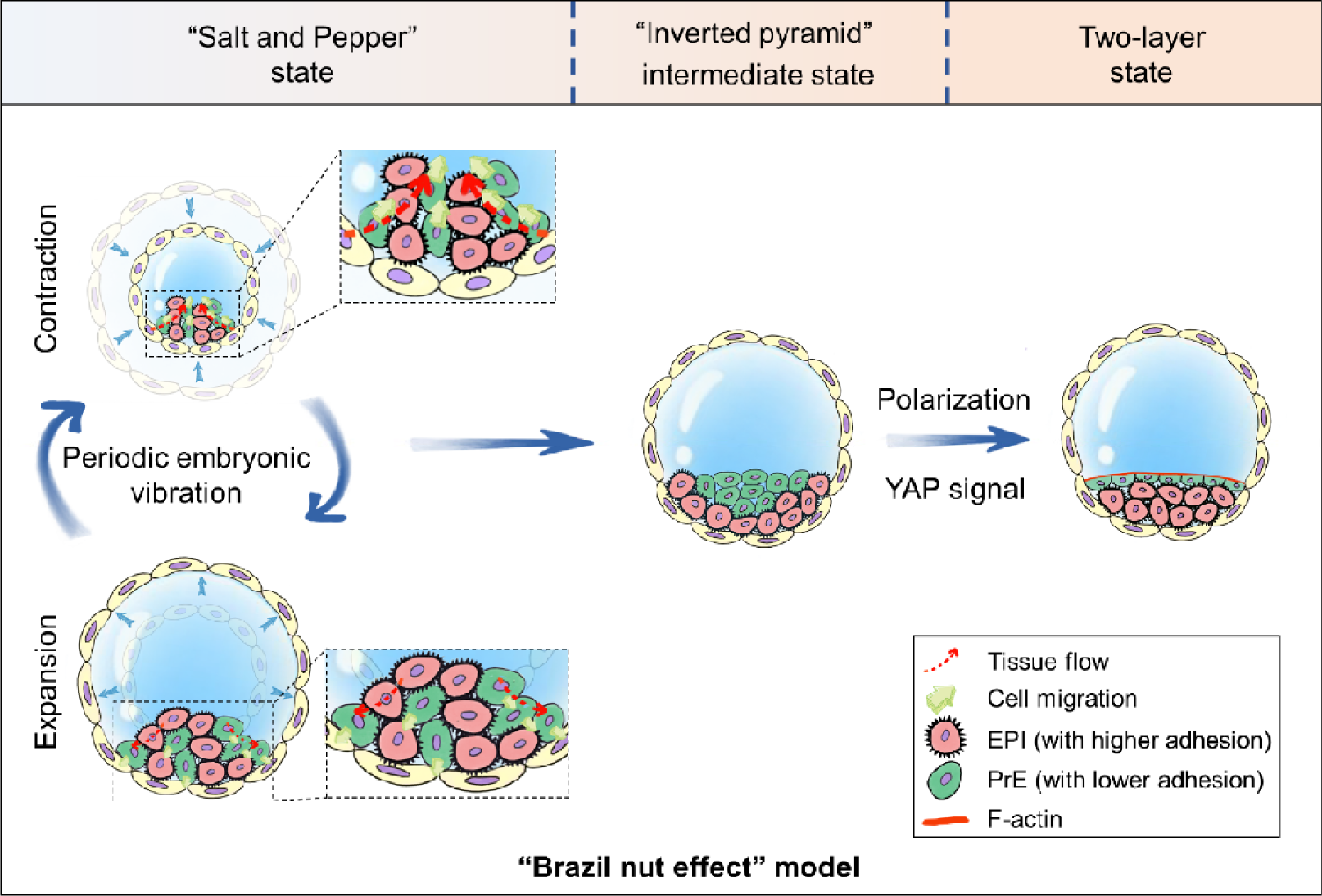
“Brazil nut effect” model of the ICM segregation. The model for the topographical regionalization of the ICM during blastocyst development in which the asymmetric tissue flows inside the ICM compartment generated by the periodic contraction-expansion of the embryo induces PrE precursors moving toward the center of the ICM/cavity boundary to form an “inverted pyramid” through viscous segregation. Then, the lateral tissue flows at the surface of the ICM during the expansion phase together with chemical signal drives PrE cells to rearrange as a monolayer epithelium to form a two-layer pattern. Blastocyst cavity vibration promotes the spatial segregation of PrE/EPI cells through “Brazil nut effect” mechanism.

It is generally accompanied by the appearance of the cavity in embryonic development and organogenesis. The appearance and mechanical behavior of cavity accompanies with the generation and accumulation of mechanical energy. When the energy of the cavity has accumulated to a certain extent, it may be released and converted in other forms, which affects the entire system. The function of cavity expansion in the lineage specification of PrE/EPI cells has also been observed in a recent study ^21^. In our experiments, drug treatment of blastocysts has certain limitations in terms of gene expression, but we are more concerned about changes in physical factors. In our results, we found that cell movement is reduced in the absence of cavity vibration without affecting cavity expansion (Figure 2e-g). However, cavity vibration depends on the tension generated by cavity expansion. Thus, these results indicate that cavity expansion is an essential requirement for cavity vibration induced pattern formation. From this point of view, the function of cavity mechanics is conservative and quite stable. In this case, the accumulation of mechanical force in the blastocyst cavity is transmitted into the movement of cells in the ICM. Similar function of tissue-scale wave of mechanochemical propagation in embryonic development has also been observed in other species ^47^.

Our results are the first to propose that in early mammalian embryonic development without a specific morphogen, the expansion of the cavity converts chemical energy into mechanical energy and stores it, which is then released by vibration and promotes the migration of cells. This cell sorting mode driven by the global geometry and mechanical behavior is very conservative and quite stable for ensuring the robustness of cell segregation and pattern formation in the development of embryos and organs. It is also helpful to guide the construction of artificial organizations for research or medical applications.

## ACKNOWLEDGMENTS

Thanks to Professor Philippe Soriano of Icahn School of Medicine at Mount Sinai for donating *Pdgfr*α*^H2B-GFP/+^* mouse to us. This work was supported by the National Key R&D Program of China (2017YFA0506500, 2016YFC1102203, and 2016YFC1101100), the National Natural Science Foundation of China (31370018, 11972206, 11902114, 11421202, 11827803, 11902020, 12072350, and 11832017), the CAS Key Research Program of Frontier Sciences (QYZDB-SSW-JSC036), the CAS Strategic Priority Research Program (XDB22040403), and Fundamental Research Funds for the Central Universities (ZG140S1971).

## AUTHOR CONTRIBUTIONS

Y.F. and J.D. designed the study and interpreted experiments. Z.G., J.C., and S.G. performed experiments. Y.J. and X.Z. did the mechanical modeling. D.Q., Z.G., L.L., J.Z., and Z.C. helped with the embryonic experiments. M.T., B.W., and F.M. helped with the statistical analysis of data. Y.F. and J.D. conceived and supervised this project and wrote the paper.

## COMPETING INTERESTS

There are no competing interests

## SUPPLEMENTARY MATERIALS

### Supplementary Figures

**Supplementary Figure 1.**
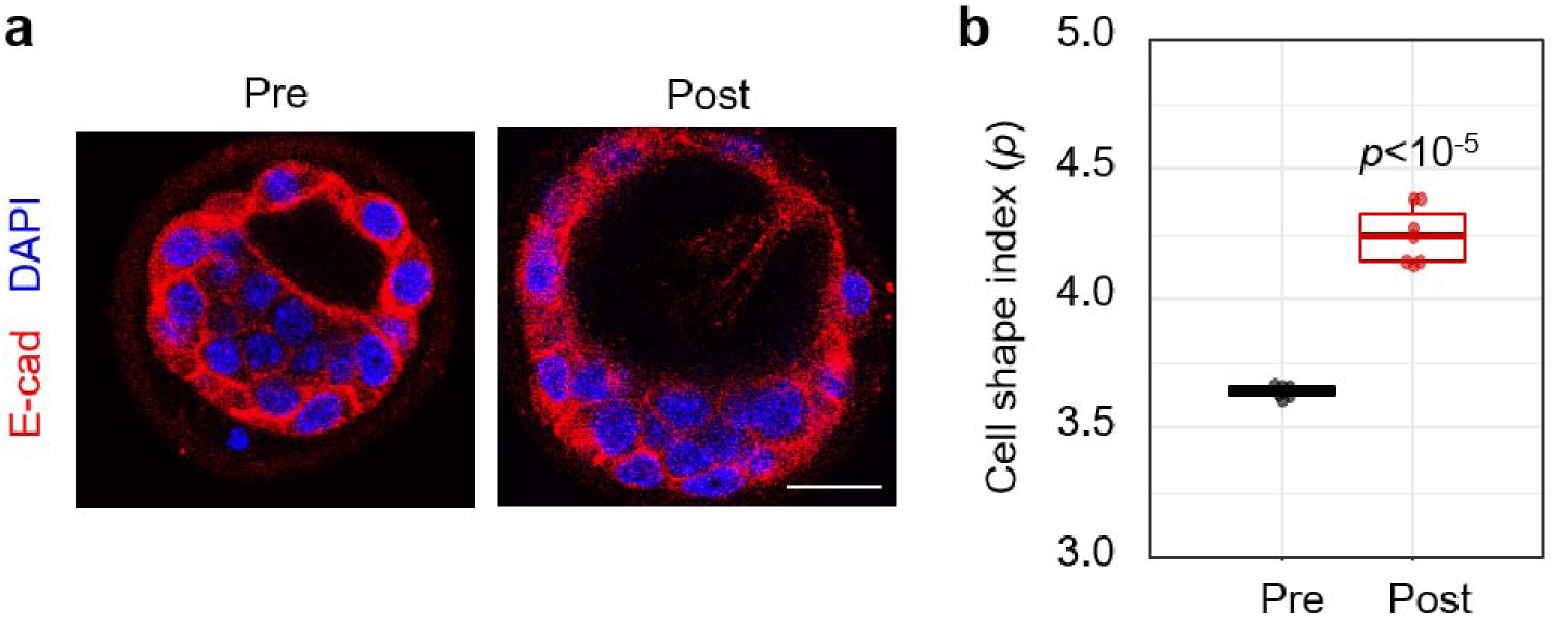
The shape index of ICM cells prior and post the emergence of rhythmic cavity vibration. (a) Representative immunofluorescence images of embryos prior and post the emergence of rhythmic cavity vibration. Cell shape is indicated by E-cadherin (E-cad) labeling. Scale bar, 25 μm. (b) The shape index of cells in ICM pre (n = 6 embryos) and post (n = 7 embryos) the emergence of rhythmic cavity vibration. The dot represents the mean value of the shape index of the cells in the ICM of an embryo. The experiments were repeated for at least three times.

**Supplementary Figure 2.**
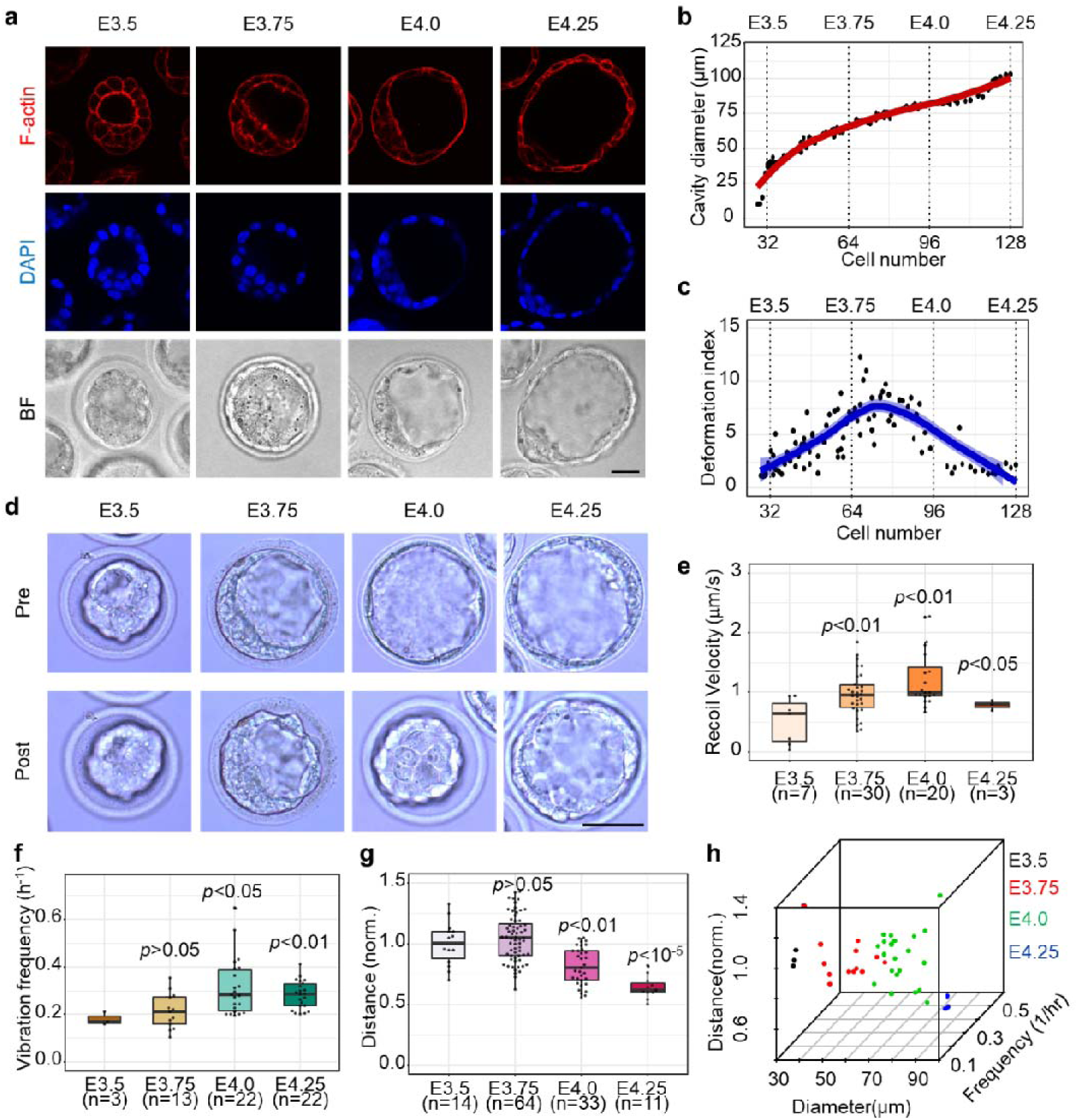
Migration of PrE precursors is correlated with tension and vibration of blastocyst cavity at different stages. (a) Representative immunofluorescence and bright field images of blastocysts at different stages. F-actin is stained by phalloidin. Scale bar, 25 μm. (b) Quantification of the cavity diameter of embryos with different cell numbers. The dots indicate the cavity diameter of an embryo. The line represents the local linear regression fitting curve of the cavity diameter of multiple embryos. N = 99 embryos. (c) Quantification of the TE cells deformation index (aspect ratio) of embryos with different cell numbers. The dots indicate the TE cells deformation index of an embryo. The line represents the local linear regression fitting curve of the TE cells deformation index of multiple embryos. n = 99 embryos. (d) The images of embryos at different stages indicating the cavity recoil post UV laser cutting. Scale bar, 50 μm. (e) The cavity recoil speed after UV laser cut of embryos at different stages. (f) The vibration frequency of the blastocyst cavity of embryos at different stages. (g) The distance of the Pdgfrα-GFP positive cells to the center of the ICM surface in embryos at different stages. The distance from the Pdgfrα-GFP positive cell at E3.5 to the center of the ICM surface is normalized as 1 (h) The correlation of cavity diameter, vibration frequency and the distance of Pdgfrα-GFP positive cells to the center of the ICM surface. n = 41 embryos. The experiments were repeated for at least three times.

**Supplementary Figure 3.**
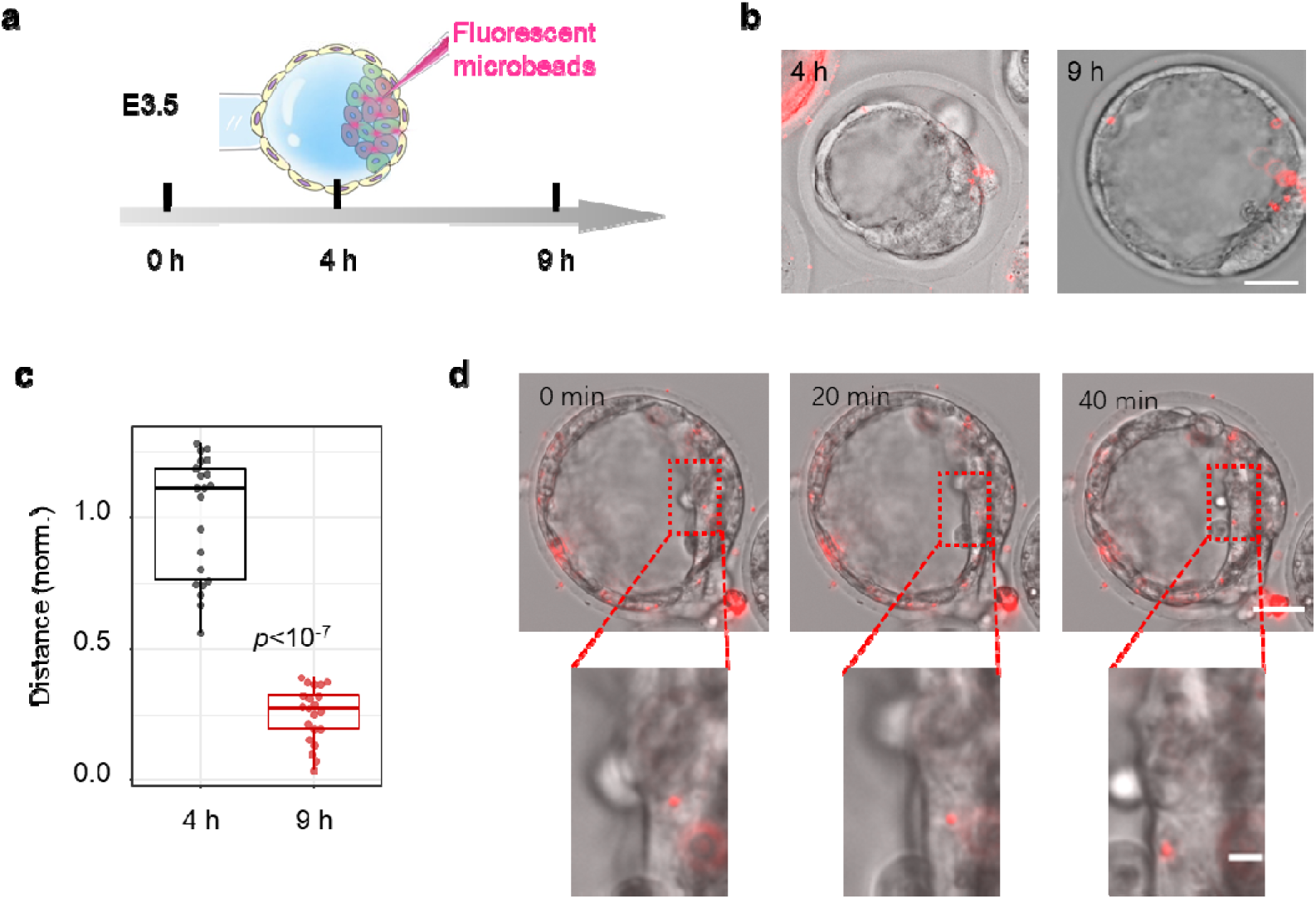
Movement of fluorescent microbeads in the ICM of the blastocyst. (a) Schematic diagram of fluorescent microbeads injection into ICM compartment at E3.5. (b) Representative images of 0 h and 5 h after injection of fluorescent beads in E3.5 embryos. Scale bar, 25 μm. (c) The distance of the fluorescent beads to the center of the ICM surface (n = 21 fluorescent beads from 7 embryos). The distance from the fluorescent microbeads at 4 h to the center of the ICM surface is normalized as 1. (d) Representative time-lapse images of fluorescent beads movement in ICM. The bottom images are enlarged views of the red box areas of the top images. Scale bar, top: 25 μm, bottom: 5 μm.

**Supplementary Figure 4.**
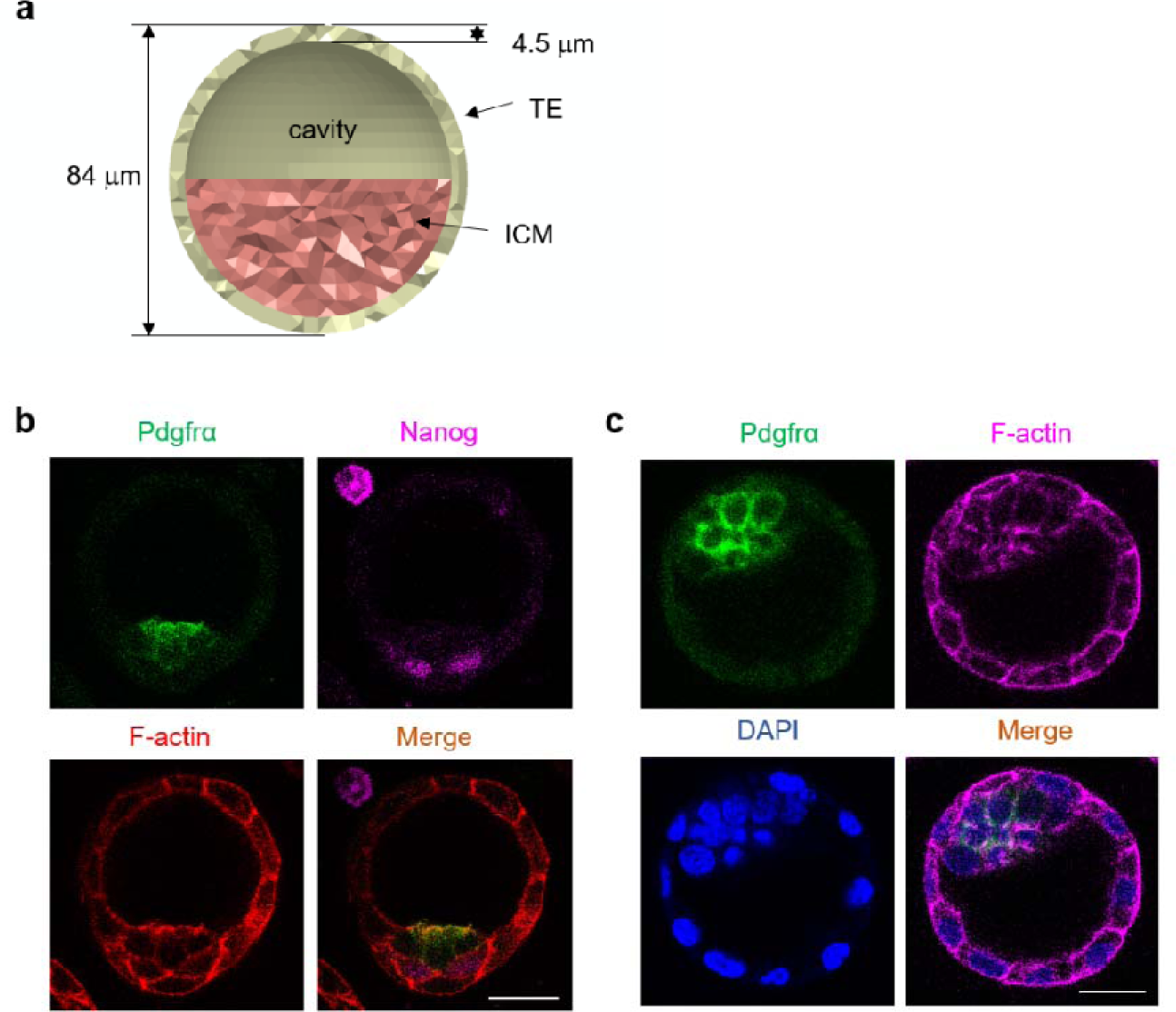
Mechanical model and F-actin assembly in blastocysts. (a) The cross-section of the finite element model in Figure 4e. (b) Representative immunofluorescence images of embryos with the Pdgfrα-GFP positive cells showing “inverted pyramid” distribution. Scale bar, 25 μm. (c) Representative immunofluorescence images of embryos with the outermost Pdgfrα-GFP positive cells showing a “Rose-like” distribution at the surface of ICM. Scale bar, 25 μm. The experiments were repeated for at least three times.

**Supplementary Figure 5.**
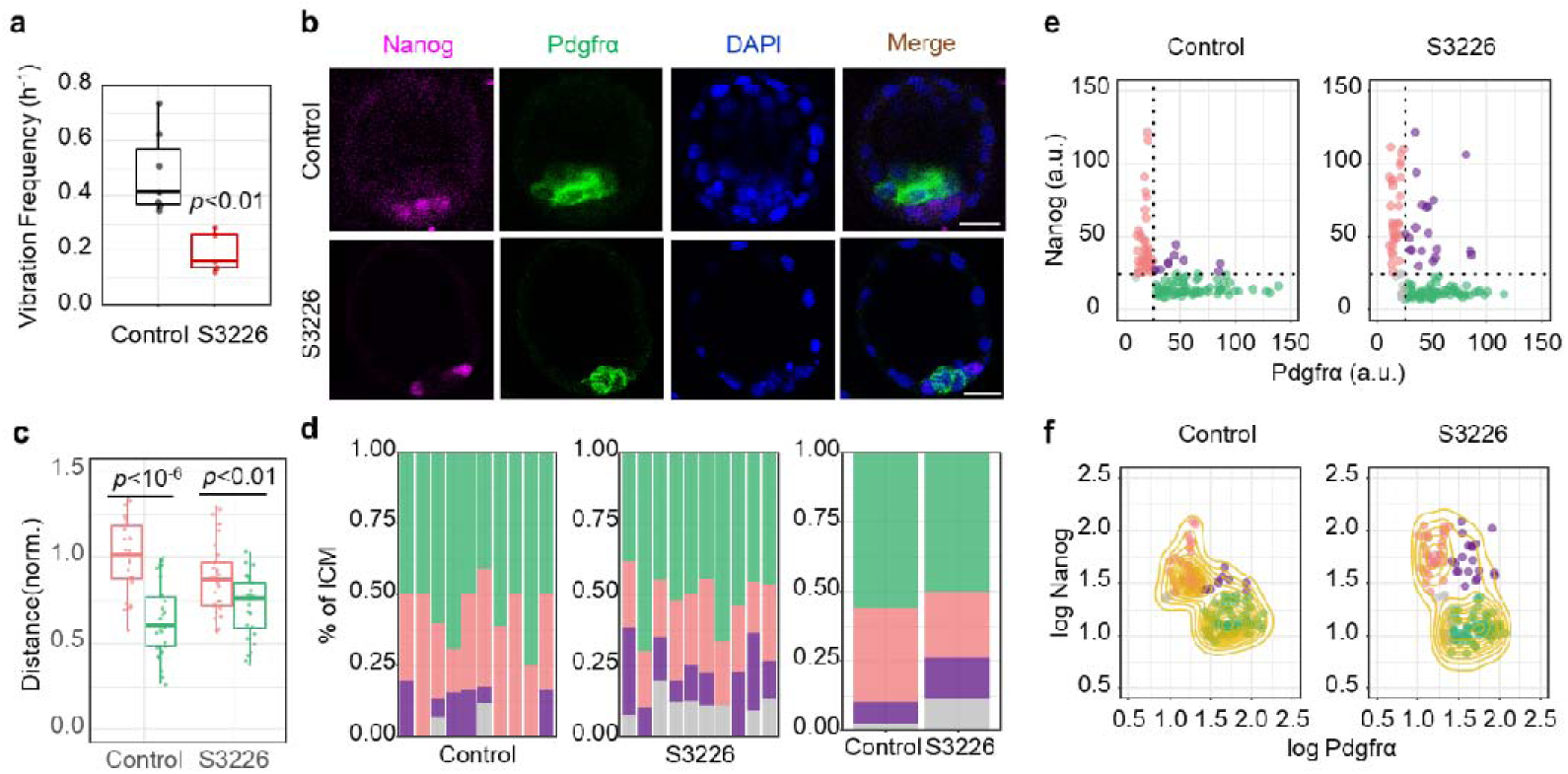
S3226 treatment inhibits the spatial segregation and lineage specification of PrE/EPI. (a) Vibration frequency of blastocyst cavity in control (n = 7 embryos) and S3226-treated (S3226, n = 5 embryos) embryos. (b) Representative immunofluorescence images of the E3.5 embryos fixed after 24 hours of S3226 treatment. Scale bar, 25 μm. (c) The distance of Nanog and Pdgfrα positive cells to the center of the ICM surface in control (Nanog: n = 23 cells from 7 embryos, Pdgfrα: n = 27 cells from 7 embryos) and S3226 treated (Nanog: n = 29 cells from 7 embryos, Pdgfrα: n = 29 cells from 7 embryos) embryos. The mean value of the distance from the nanog cells of the control group to the center of the ICM surface is normalized as 1. (d) Average ICM composition at the end of the culture period for embryos treated with S3226, shown as % of the ICM. Control: n = 129 cells from 10 embryos. S3226: n = 133 cells from 10 embryos. (e) Scatter plot of fluorescence intensity levels of Nanog and Pdgfrα after S3226 treatment. DN, gray, double negative (Nanog-, Pdgfrα-); DP, purple, double positive (Nanog +, Pdgfrα+); EPI, red, (Nanog +, Pdgfrα-); PrE, green, (Nanog-, Pdgfrα-). (f) Scatter plots for same data as in (e), represented as logarithm. The yellow contour lines show the density. The experiments were repeated for at least three times.

**Supplementary Figure 6.**
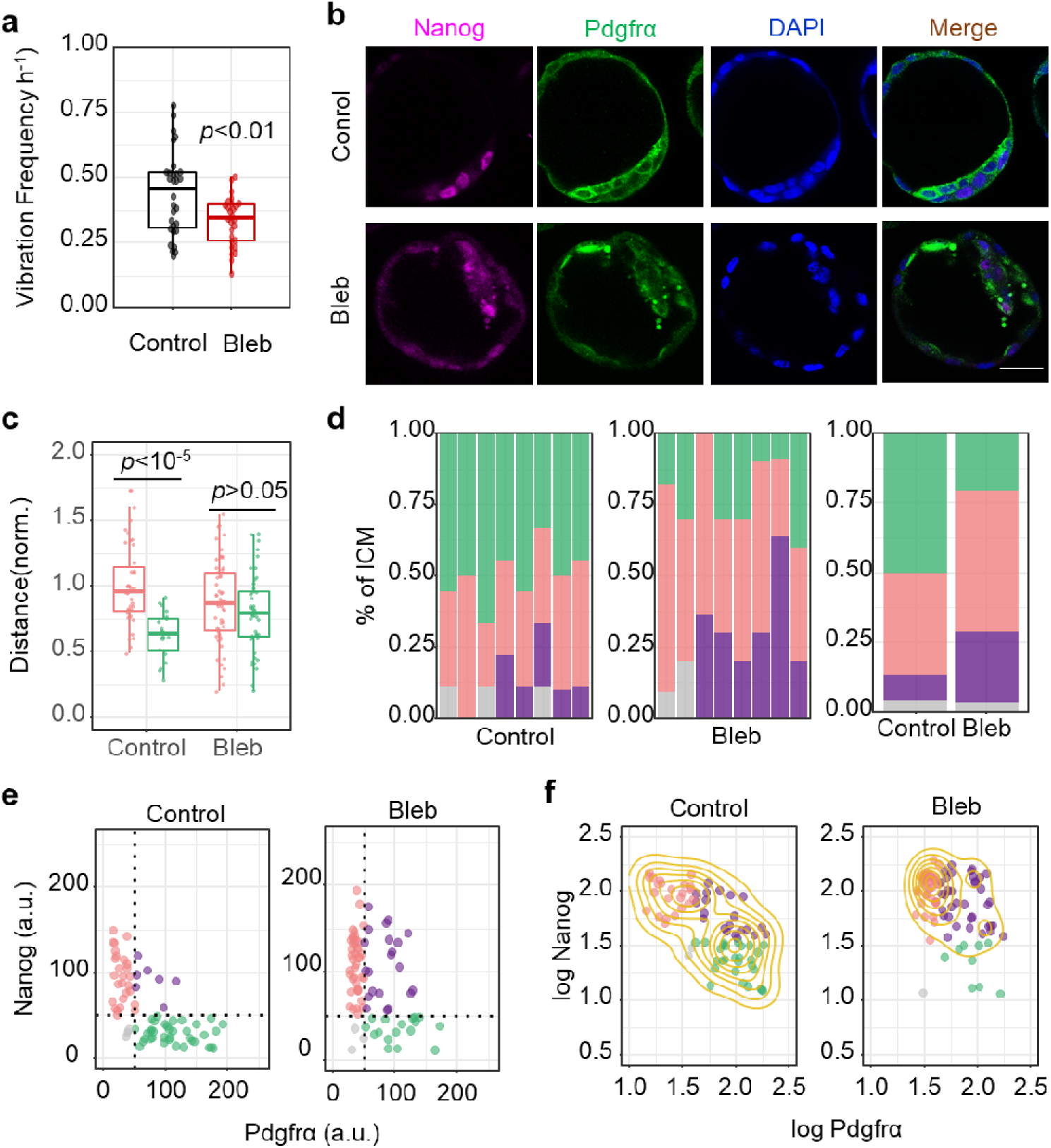
Blebbistatin treatment inhibits the spatial segregation and lineage specification of PrE/EPI. (a) Vibration frequency of blastocyst cavity in control (n = 30 embryos) and Blebbistatin-treated (Bleb, n = 27 embryos) embryos. (b) Representative immunofluorescence images of the E3.5 embryos fixed after 24 hours of Bleb treatment. Scale bar, 25 μm. (c) The distance of Nanog and Pdfrα positive cells to the center of the ICM surface in control (Nanog: n = 39 cells from 7 embryos, Pdgfrα: n = 21 cells from 7 embryos) and Bleb treated (Nanog: n = 53 cells from 6 embryos, PDGFRα: n = 46 cells from 6 embryos) embryos. The mean value of the distance from the nanog cells of the control group to the center of the ICM surface is normalized as 1. (d) Average ICM composition at the end of the culture period for embryos treated with Bleb, shown as % of the ICM. Comtrol: n = 74 cells from 8 embryos. Bleb: n = 83 cells from 8 embryos. (e) Scatter plot of fluorescence intensity levels of Nanog and Pdgfrα after Bleb treatment. DN, gray, double negative (Nanog-, Pdgfrα-); DP, purple, double positive (Nanog +, Pdgfrα+); EPI, red, (Nanog +, Pdgfrα-); PrE, green, (Nanog-, Pdgfrα-). (f) Scatter plots for same data as in (e), represented as logarithm. The yellow contour lines show the density. The experiments were repeated for at least three times.

**Supplementary Figure 7.**
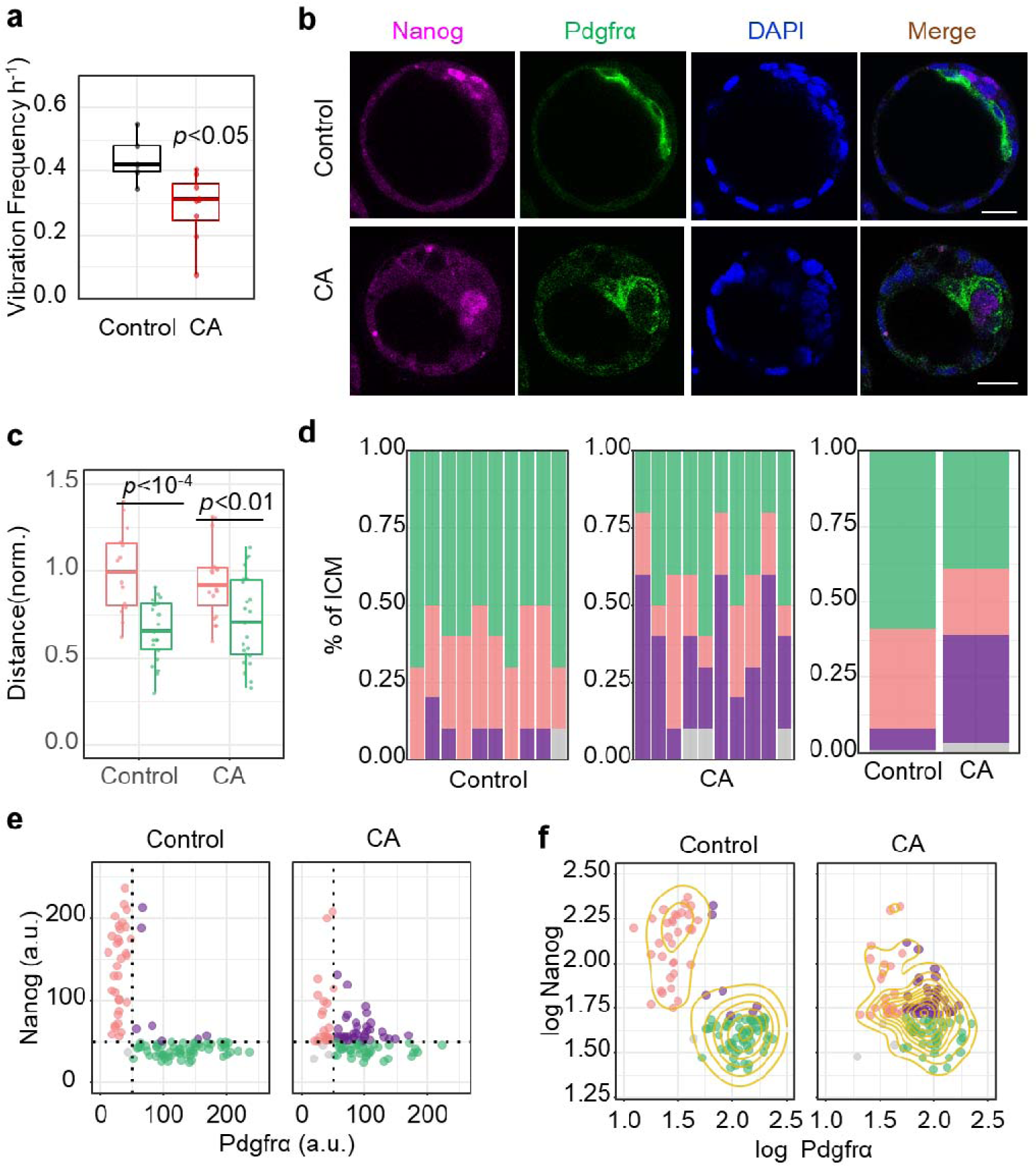
Cytochalasin A treatment inhibits the spatial segregation and lineage specification of PrE/EPI layers. (a) Vibration frequency of blastocyst cavity in control (Control, n = 5 embryos) and cytochalasin A treatment (CA, n = 8 embryos). (b) Representative immunofluorescence images of the E3.5 embryos fixed after 24 hours of CA treatment. Scale bar, 25 μm. (c) The distance of Nanog and Pdfrα positive cells to the center of the ICM surface in control (Nanog: n = 15 cells from 5 embryos, Pdgfrα: n = 21 cells from 5 embryos) and CA treated (Nanog: n = 19 cells from 5 embryos, Pdgfrα: n = 23 cells from 5 embryos) embryos. The mean value of the distance from the nanog cells of the control group to the center of the ICM surface is normalized as 1. (d) Average ICM composition at the end of the culture period for embryos treated with CA, shown as % of the ICM. Comtrol: n = 100 cells from 10 embryos. CA: n = 100 cells from 10 embryos. (e) Scatter plot of fluorescence intensity levels of Nanog and Pdgfrα after CA treatment. DN, gray, double negative (Nanog-, Pdgfrα-); DP, purple, double positive (Nanog+, Pdgfrα+); EPI, red, (Nanog+, Pdgfrα-); PrE, green, (Nanog-, Pdgfrα-). (f) Scatter plots for same data as in (e), represented as logarithm. The yellow contour lines show the density. The experiments were repeated for at least three times.

**Supplementary Figure 8.**
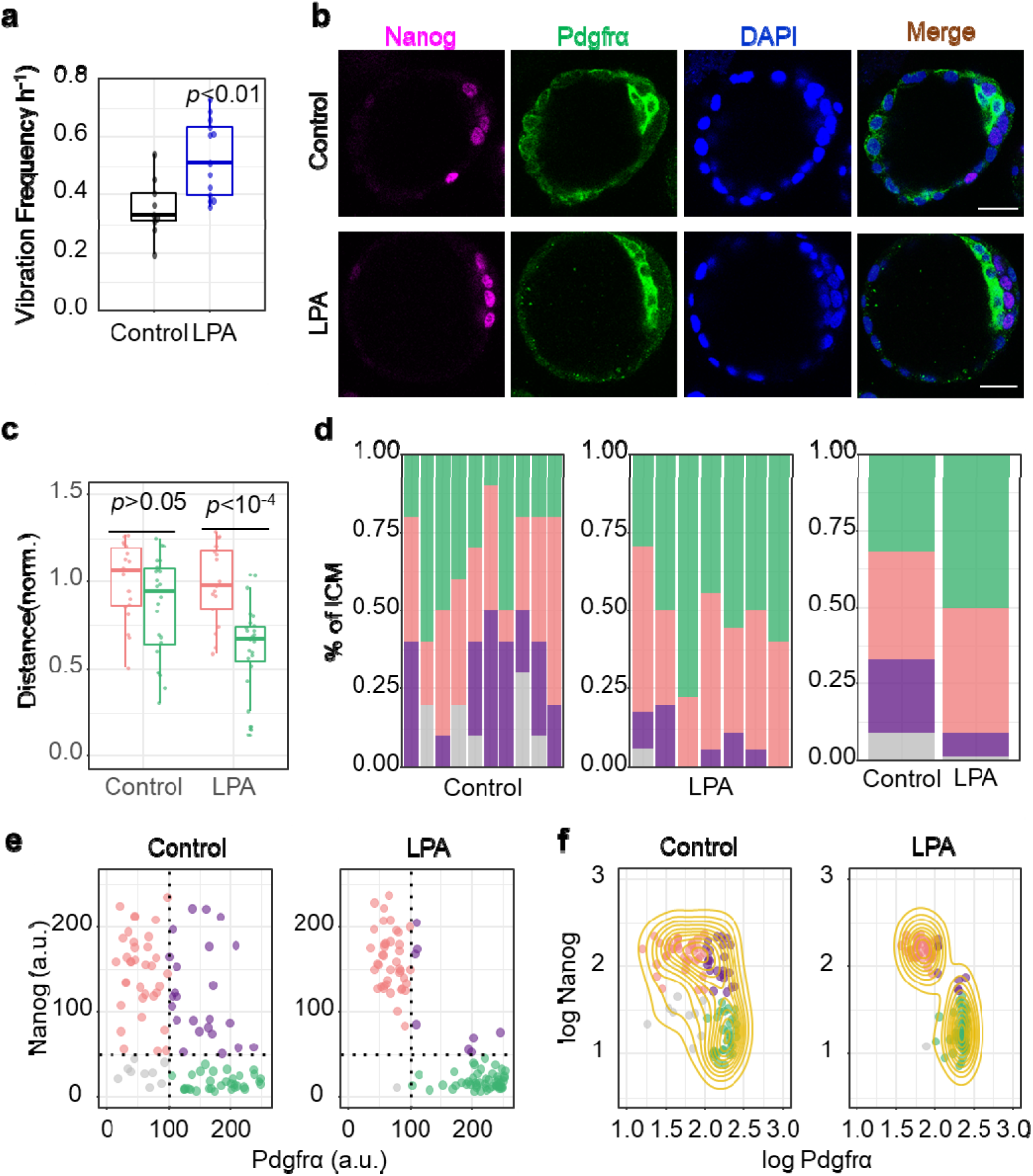
LPA treatment facilitates the spatial segregation and lineage specification of PrE/EPI. (a) Vibration frequency of blastocyst cavity in control (n = 9 embryos) and LPA treatment (LPA, n = 13 embryos). (b) Representative immunofluorescence images of the embryos fixed after 24 hours of LPA treatment. Scale bar, 25 μm. (c) The distance of Nanog and Pdgfrα positive cells to the center of the ICM surface in control (Nanog: n = 18 cells from 5 embryos, Pdgfrα: n = 24 cells from 5 embryos) and LPA-treated (Nanog: n = 17 cells from 5 embryos, Pdgfrα: n = 23 cells from 5 embryos) embryos. The mean value of the distance from the nanog cells of the control group to the center of the ICM surface is normalized as 1. (d) Average ICM composition at the end of the culture period for embryos treated with LPA, shown as % of the ICM. Comtrol: n = 100 cells from 10 embryos. LPA: n = 100 cells from 7 embryos. (e) Scatter plot of fluorescence intensity levels of Nanog and Pdgfrα after LPA treatment. DN, gray, double negative (Nanog-, Pdgfrα-); DP, purple, double positive (Nanog+, Pdgfrα+); EPI, red, (Nanog +, Pdgfrα-); PrE, green, (Nanog-, Pdgfrα-). (f) Scatter plots for same data as in (e), represented as logarithm. The yellow contour lines show the density. The experiments were repeated for at least three times.

**Supplementary Figure 9.**
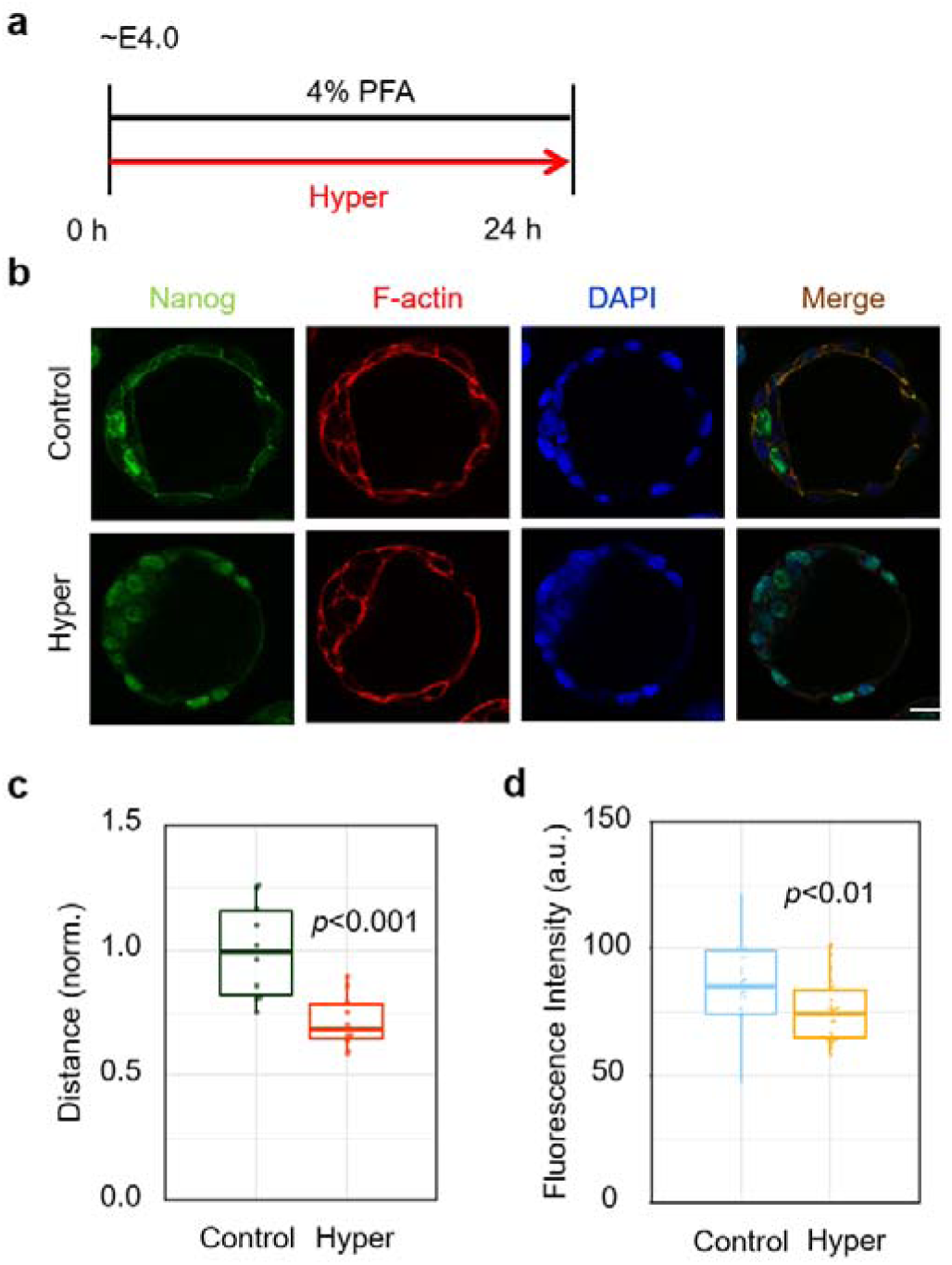
Hypertonic treatment reverses the spatial segregation and lineage specification of PrE/EPI. (a) Schematic diagram of the process of hypertonic treatment of blastocysts. The length of the arrow represents the duration of embryo culture. (b) Representative immunofluorescence images of embryos treated as indicated in (a). Scale bar, 25 μm. (c) The distance of the Nanog positive cells to the center of the ICM surface. n = 10 cells from 2 embryos. The mean value of the distance from the nanog cells of the control group to the center of the ICM surface is normalized as 1. (d) The fluorescence intensity of Nanog positive cells in control and hypertonicity-treated embryos. n = 30 cells from 6 embryos. The experiments were repeated for at least three times.

**Supplementary Figure 10.**
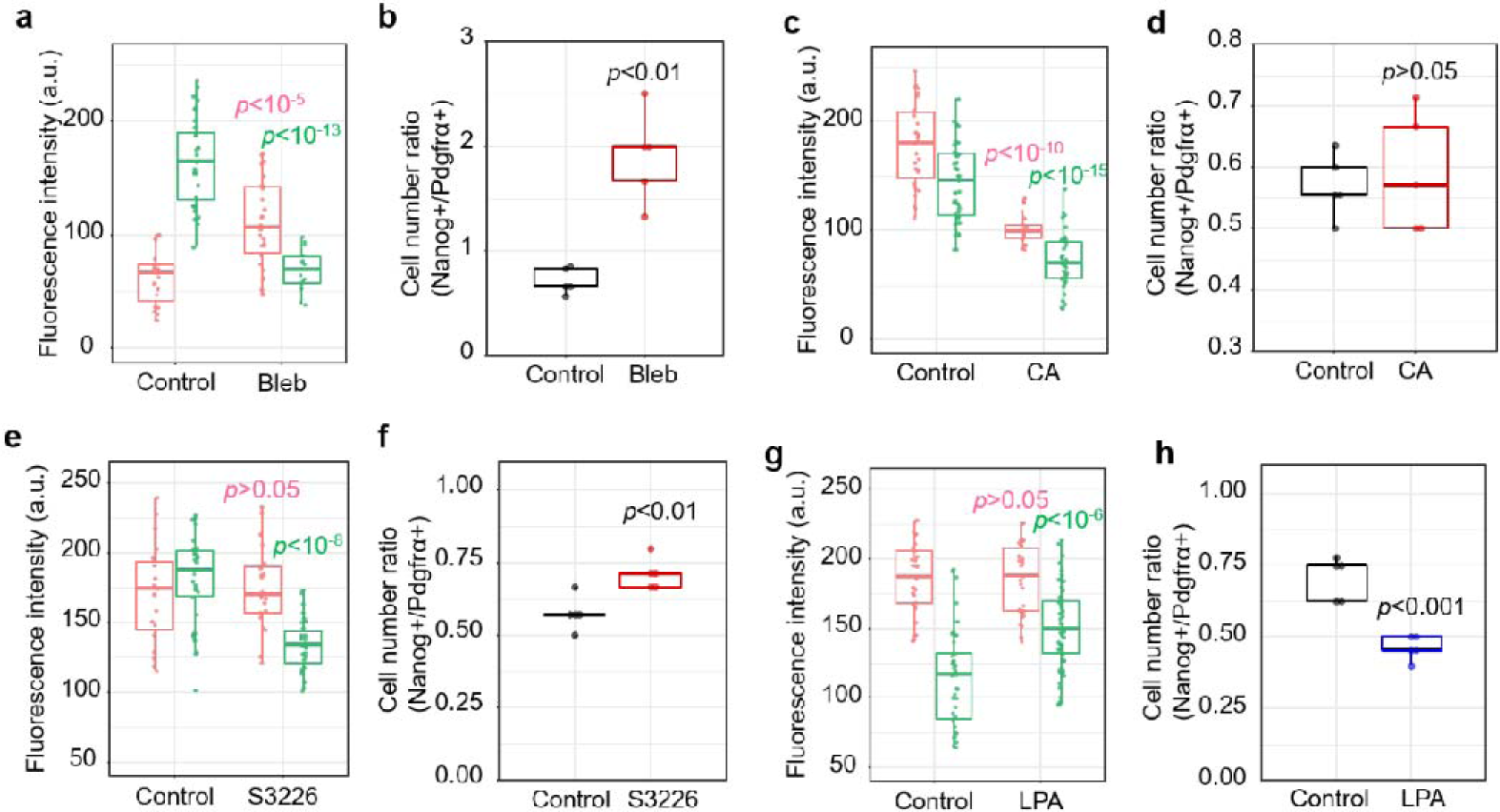
Gene expression pattern of ICM cells in each treatment group (Bleb, CA, S3226, and LPA) (a) The fluorescence intensity of control (Nanog: n = 23 cells from 5 embryos, Pdgfrα: n = 32 cells from 5 embryos) and Bleb-treated (Nanog: n = 26 cells from 5 embryos, Pdgfrα: n = 14 cells from 5 embryos) embryos. (b) The cell number ratio (Nanog+/Pdgfrα+) of control and Bleb-treated embryos in ICM. The same data as in (a). (c) The fluorescence intensity of control (Nanog: n = 27 cells from 5 embryos, Pdgfrα: n = 47 cells from 5 embryos) and CA-treated (Nanog: n = 22 cells from 5 embryos, Pdgfrα: n = 37cells from 5 embryos) embryos. (d) The cell number ratio (Nanog+/Pdgfrα+) of control and CA-treated embryos in ICM. The same data as in (c). (e) The fluorescence intensity of control (Nanog: n = 19 cells from 5 embryos, Pdgfrα: n = 33 cells from 5 embryos) and S3226-treated (Nanog: n = 22 cells from 5 embryos, Pdgfrα: n = 31 cells from 5 embryos) embryos. (f) The cell number ratio (Nanog+/Pdgfrα+) of control and S3226-treated embryos in ICM. The same data as in (e). (g) The fluorescence intensity of control (Nanog: n = 29 cells from 5 embryos, Pdgfrα: n = 41cells from 5 embryos) and LPA-treated (Nanog: n = 26 cells from 5 embryos, Pdgfrα: n = 56 cells from 5 embryos) embryos. (h) The cell number ratio (Nanog +/Pdgfrα+) of control and LPA-treated embryos in ICM. The same data as in (g). The experiments were repeated for at least three times.

### Supplementary Movies

**Movie S1. Approach of PrE Cells to the Center of the ICM Surface, Related to Figure 1b**

Imaging live embryo expressing *Pdgfr*α*^H2B-GFP/+^* shows that the relative allocation of Pare cells tended to approach to the center of ICM surface after E3.75. The scale bar is 20 μm. Time is indicated in h.

**Movie S2. Rhythmic Vibration of the Blastocyst Cavity, Related to Figure 1f**

Live-imaging of an embryo shows that the blastocyst cavity continues to expand until E3.75, after which the blastocyst begins to display rhythmic vibration. Each contraction is marked in the upper right corner of the video. The scale bar is 20 μm. Time is indicated in h.

**Movie S3. Phase Change of ICM, Related to Figure 1h**

Movie shows the heat map of the velocity field during embryonic development. At about 9h, the ICM undergoes a phase change from solid-like to liquid-like. The scale bar is 20 μm. The color bar of the heat map is 0 ∼ 9 μm/h. Time is indicated in h.

**Movie S4. Hypertonic Treatment Inhibits Cell Motility, Related to Figure 2c**

(a) Movie shows the movement of PrE cells in a live embryo expressing *pdgfr*α^H2B-GFP/+^. (b) Movie shows live embryo treated with hypertonicity, which indicates that hypertonic treatment can significantly inhibit cell movement. The scale bar is 20 μm. Time is indicated in h.

**Movie S5. Direction of the Cell Movement during the Contraction of Embryo, Related to Figure 3b**

Live-imaging of an embryo expressing *pdgfr*α*^H2B-GFP/+^* shows that the movement of PrE cells during contraction is towards the center of the ICM, demonstrating the consistency of tissue flows and the movement direction of PrE cells. The scale bar is 20 μm. Time is indicated in h.

**Movie S6. Direction of the Cell Movement during the Expansion of Embryo, Related to Figure 3d**

Live-imaging of an embryo expressing *pdgfr*α*^H2B-GFP/+^* indicates that during the expansion of blastocoel, the direction of tissue flows is paralleled with the ICM/cavity boundary. The scale bar is 20 μm. Time is indicated in h.

**Movie S7. Simulations of 10% embryo contraction and expansion with a simplified continuous model, Related to Figure 3e**

The model was to illustrate the trend of velocity vector distribution during contraction and expansion.

a. Distribution of velocity magnitude in contraction from 0-60s.
b. Distribution of velocity orientation in contraction from 0-60s.
c. Distribution of velocity magnitude in expansion from 0-3600s.
d. Distribution of velocity orientation in expansion from 0-3600s.

**Movie S8. Movement of Microbeads after Embryo Contraction, Related to Supplementary Figure s3d**

Imaging live embryo injected with fluorescent microbeads shows the movement of microbeads after embryo contraction. The scale bar is 20 μm. Time is indicated in min.

**Movie S9. Simulations of cell viscous segregation during cavity vibration with a simplified finite element model, Related to Figure 3i**

The model was to illustrate the effect of cohesive properties and periodic contraction-expansion of embryos on the separation of PrE and EPI cells.

a. The separation process of PrE and EPI cells with different cohesive properties (EPI: red, strong cohesiveness; PrE: blue, week cohesiveness) under periodic embryo contraction-expansion.
b. The separation failed when cohesive properties of cells were the same.
c. The separation failed when embryo contraction speed reduced to one-third.

**Movie S10. Fluorescence intensity alteration of *pdgfrα^H2B-GFP/+^* during development, Related to Figure 7a**

Live-imaging of an embryo expressing *pdgfr*α*^H2B-GFP/+^* indicates that during the migration of Pdgfra positive cells, GFP expression intensity gradually increased. The scale bar is 20 μm. Time is indicated in h.

### Mechanical Modeling

To assess the motion pattern of the embryo compound during contraction and expansion, a simplified continuous model was developed. As shown in Supplementary Fig. S3a, the model consists of TE, ICM, and the cavity. The geometric information of the model was based on the microscope images of the embryo. The embryo was assumed to be isotropic material with the modulus of 360 Pa, Poisson ratio of 0.45, and density of 1000 kg/m^3^. The model was meshed with 4-node linear tetrahedron in the approximate size of 4 μm. A contraction of 10% diameter in 2 minutes and an expansion of 10% diameters in 2 hours was simulated. The distributions of velocity magnitude and orientation were shown in Fig. 3e.

To study the effect of cohesive properties and periodic contraction-expansion of embryo on the separation of PrE and EPI cells, a simplified finite element model of embryo was developed (Figure 3i). The geometric information of the model was based on the microscope images of the embryo. The embryo was simplified as a sphere with the diameter of 84 μm. PrE and EPI cells were simplified as spheres with the diameter of 13 μm. Material property of all cells was assumed to be isotropic with the elastic modulus of 360 Pa, Poisson ratio of 0.45, and density of 1000 kg/m^3^. A liquid damping coefficient of 9.4e^-11^ MPa*s was applied. The model was meshed with 4-node linear tetrahedron in the approximate size of 5 μm. To accelerate the simulation, the periods of contraction and expansion was set to 0.36 s and 0.54 s, and a mass scaling acceleration algorithm was applied. 7 cycles were simulated. Since the real expansion was a quasi-static process, the kinetic energy was set to 0 at the beginning of each cycle. The cohesive properties between PrE, EPI, and TE cells were assumed to be different (Table 1). To study the effect of different cohesive properties on the separation of PrE and EPI cells, a control case with the same cohesive property (= Y-Z cohesive property) was developed. To study the effect of different contraction speed on the separation of PrE and EPI cells, a control case with the contraction speed 3 times slower was developed (Figure 3i).

**Table 1:**
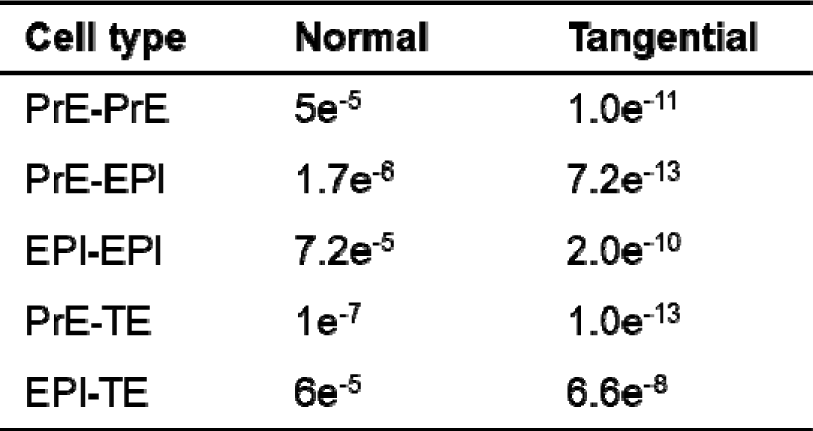
Cohesive stiffness coefficient between PrE, EPI, and TE cells.

### MATERIALS AND METHODS

#### Mouse Husbandry

All mice were bred and reared in the animal facility of Tsinghua University at 22 °C with a 12-hour light/dark cycle (lighting time 7:00-19:00). Food and water are freely available. All animal studies were conducted under the guidance of the Animal Care and Utilization Committee (IACUC) of Tsinghua University. According to the National Institutes of Health “Animal Ethical Use Guidelines”, the experimental procedure has been approved by the Laboratory Animal Care and Use Management Committee of Tsinghua University and the Beijing Municipal Science and Technology Commission (SYXK-2019-0044). The *Pdgfr*α*^H2B-GFP/+^* mouse were donated by Professor Philippe Soriano of Icahn School of Medicine at Mount Sinai.

#### Embryo Collection and Culture

In order to obtain preimplantation embryos, female mice (2~3 months old) were injected intraperitoneally with 10 international units (IU) of pregnant mare serum gonadotropin (PMSG)(Solarbio,P9970), and then injected with 10 international units (IU) of human chorionic gonadotropin (hCG)(ShuSheng,11025) 48 hours later. After the injection of hCG, the female mice in the superovulation period were directly mated with male mice (3~4 months old). The pre-implanted embryo was flushed out of the uterus or fallopian tube with KSOM(Caisson,IVL04). Then embryos were cultured in drops of KSOM under mineral oil (Sigma,M8410) at 37 0 in a 5% CO_2_ atmosphere. Stereo microscope for embryo manipulation: Olympus SZX16.

#### Pharmacological Treatment

Sucrose(Macklin, S818046) was dissolved in ddH_2_O, the concentration of the stock solution was 1 mol/L. The stock solution is diluted by KSOM (260 mOsm) into a hypertonic solution(340 mOsm) of sucrose with a final concentration of 100 mM. We used ddH_2_O to dilute KSOM into a 70% hypotonic solution (170 mOsm). (-)-Blebbistatin (Sigma,B0560) was dissolved in DMSO at a concentration of 25 mM. The stock solution was diluted by KSOM to a working concentration of 1 μM. Dissolve LPA(Cayman,62215) in ethonal at the stock concentration of 1 mM. The stock solution was diluted by KSOM to a working concentration of 5 μM. Cytochalasin A(Sigma,C6637) was dissolved in DMSO at the stock concentration of 0.5 mg/mL. The stock solution was diluted by KSOM to a working concentration of 1 μg/mL. Dissolved S3226(Sigma,SML 1996) in ddH_2_O to prepare a stock solution with a concentration of 2 mg/mL. The stock solution was diluted by KSOM to a working concentration of 20 nM. Dissolved Verteporfin(Tocris,5305) in DMSO to prepare a stock solution with a concentration of 10 mg/mL. The stock solution was diluted by KSOM to a working concentration of 2 μg/mL. Dissolved Aphidicolin (Abcam,ab142400) in DMSO to prepare a stock solution with a concentration of 10 mg/mL. The stock solution was diluted by KSOM to a working concentration of 2 μg/mL.

#### Live Embryo Imaging

For real-time imaging, embryos were cultured in 20 μL of KSOM drops covered with 2 mL of mineral oil on a 35 mm glass bottom dish in an environmental chamber at 37 °C with 5% CO_2_. Live embryo image: Andor Dragonfly, Leica DMi8(leica application suite X 2.0).

#### Embryo immunofluorescence staining

Embryos were washed in Dulbecco’s phosphate buffered saline (DPBS)(Corning,21-031-CVR) containing 1% (w/v) polyvinylpyrrolidone (PVP)(Sigma,PVP40) and fixed with 4% PFA(Sigma,P6148) for 15 minutes at room temperature, then infiltrated with DPBS (0.5% Triton-100), placed at room temperature for 20 minutes, and then transferred to blocking buffer (DPBS containing 0.1% Tween-20 at 4°C; 3% BSA(Solarbio,A8020)) for at least 4 hours. The embryos were incubated with primary antibodies diluted in DPBS at 4°C overnight. The embryos were then washed for 3 times in DPBS-PVP, and incubated with secondary antibodies diluted in DPBS for 2 hours at room temperature. Then the embryos were washed for 3 times with DPBS-PVP, and then placed in DAPI(Solarbio,C0060) solution for 15 minutes. Finally, the embryos were washed for 3 times with DPBS-PVP for imaging. Confocal image: Leica sp8(leica application suite X 3.3),HoloMonitor M4(HStudio 2.7).

The antibody information of the immunofluorescence experiment is as follows:

Rabbit pAb to beta Catenin(Abcam,ab16051;1:200)

Mouse mAb to E Cadherin(Abcam,ab76055;1:200)

Rabbit pAb to nanog(Abcam,ab80892;1:200)

Rabbit mAb to PDGFR alpha (HuaBio,ET1702-49;1:250)

Mouse mAb to YAP1(Abnova, H00010413-Q01;1:250)

Alexa Fluor 546 donkey anti-mouse secondary antibody (Lifetechnologies,A10036;1:200)

Alexa Fluor 568 goat anti-rabbit secondary antibody (Lifetechnologies,A11011;1:200)

DyLight 488 sheep anti-mouse secondary antibody(Novus, NBP1-72924,1:500)

DyLight 488 goat anti-rabbit secondary antibody(Novus, NBP1-72944,1:500)

Anti-Mouse Nanog eFluor 660(Invitrogen,50-5761-80,1:50)

Alexa Flour 546 phalloidin(Invitrogen,A22283;1:200)

#### Fluorescent Microbeads Injection

Embryos were placed in 20 μL of KSOM drops covered with 2 mL of mineral oil on a 35 mm dish. The embryo was held by the holder of the microinjector. We used the needle of the microinjector, which is connected to a syringe to inject fluorescent polystyrene spheres (Thermo scientific) with a diameter of 1 μm into the ICM of the blastocyst.

#### UV laser cutting

The blastocyst is placed on a glass slide with 20 μL KSOM droplets. The center of the blastocyst cavity was penetrated with a UV laser by leica LMD 7000 (laser Power: 30, laser Aperture: 3, laser Speed: 13, laser head current: 83%,laser pulse frequency: 283Hz) under a 40x fold lens. At the same time, a live cell imaging system was used to record. After the laser cutting is completed, the embryos were quickly transferred to the aforementioned culture environment.

#### Fluorescence Intensity Analysis

Fluorescence intensity was measured by Fiji ImageJ. We identified every cell in the ICM based on the fluorescence images of DAPI. Through the freehand-selection option, we calculated the average Intensity of Nanog (expressed in nucleus) fluorescence channel. Then the average Intensity of Pdgfrα (expressed on cell membrane) fluorescence channel was obtained by the freehand-line option.

#### Measurement of Cell Distance to ICM Surface Center

According to images of live embryo expressing *Pdgfr*α*^H2B-GFP/+^*, we connected the geometric center of the ICM and the geometric center of the cavity to obtain the symmetry axis of the embryo. The intersection of the symmetry axis of the embryo and the boundary of the ICM near the cavity was regarded as the center of the ICM. Then we can calculate the distance from the center of cells expressing *Pdgfr*α*^H2B-GFP/+^* to the center of the ICM.

#### PIV (Particle Image Velocimetry) Measurement

PIV analysis was performed using a custom algorithm based on MATLAB’s PIVlab software package. We use live cell image sequences of embryos to analyze the tissue flows of ICM during the contraction and expansion of the blastocoel. The average speed is subtracted from calculated velocity fields to avoid the influence of embryo movement on the calculation results. The heat map of ICM’s magnitude was also exported after being subtracted the mean velocity.

#### Shape Index and Cell Tracking

In order to determine the cell boundaries, we used a semi-automatic segmentation pipeline. The cell boundaries of ICM were obtained from the immunofluorescence images of embryos marked with ECA, and then we obtained the perimeter and area. The shape index p=P/√A, where P and A are the cell perimeter and projected area. We used the software Bitplane Imaris 9.0.1 to get the cell tracking images.

#### Significant Difference Analysis

Statistical analysis was performed using Excel and R. The continuous quantitative data were analyzed by the normality test first and then compared with the t-test or the Wilcoxon signed-rank test, and p<0.05 (two-tailed) was considered as statistically significant.

## Notes

### Competing Interest Statement

The authors have declared no competing interest.

